# Shifting roles of *Drosophila* pair-rule gene orthologs: segmental expression and function in the milkweed bug *Oncopeltus fasciatus*

**DOI:** 10.1101/721217

**Authors:** Katie Reding, Mengyao Chen, Yong Lu, Alys M. Cheatle Jarvela, Leslie Pick

## Abstract

The discovery of pair-rule genes (PRGs) in *Drosophila* revealed the existence of an underlying two-segment-wide prepattern directing embryogenesis. The milkweed bug *Oncopeltus*, a hemimetabolous insect, is a more representative arthropod: most of its segments form sequentially after gastrulation. Here we report the expression and function of orthologs of the complete set of nine *Drosophila* PRGs in *Oncopeltus*. Seven *Of*-PRG-orthologs are expressed in stripes in the primordia of every segment, rather than every-other segment, *Of-runt* is PR-like, and several are also expressed in the segment addition zone. RNAi-mediated knockdown of *Of-odd-skipped*, *paired* and *sloppy-paired* impacted all segments, with no indication of PR-like register. We confirm that *Of*-*E75A* is expressed in PR-like stripes, although it is not PR in *Drosophila*, demonstrating the existence of an underlying PR-like prepattern in *Oncopeltus*. These findings reveal that a switch occurred in regulatory circuits leading to segment formation: while several holometabolous insects are “*Drosophila*-like,” utilizing PRG-orthologs for PR-patterning, most *Of-*PRGs are expressed segmentally in *Oncopeltus*, a more basally-branching insect. Thus, an evolutionarily stable phenotype – segment formation – is directed by alternate regulatory pathways in diverse species.

**Summary Statement:** Despite the broad of conservation of segmentation in insects, the regulatory genes underlying this process in *Drosophila* have different roles in the hemipteran, *Oncopeltus fasciatus*.

## Introduction

Mechanisms directing the formation of the basic segmented body plan have been unraveled for the model insect, *Drosophila melanogaster* (reviewed in (Wieschaus and Nüsslein-Volhard, 2016). This study identified a set of pair-rule mutants, characterized by absence of alternate body segments, revealing that patterning of single segments is preceded by pre-patterning of a double-segment-wide unit that is repeated along the anterior-posterior axis of the embryo at half the frequency of segment number. Most of the pair-rule genes (PRGs) responsible for this pre-pattern are expressed in seven stripes in the *Drosophila* blastoderm, with PRG expression foreshadowing the corresponding mutant phenotype for individual PRGs (Pair-rule stripes, Figure 1). For example, *even-skipped (eve)* and *fushi tarazu (ftz)* are expressed in complementary seven-stripe patterns, each in the primordia of the alternate parasegments missing in *eve* or *ftz* mutants (Lawrence and Johnston, 1989). Other PRGs are expressed in similar complementary patterns, with the combined, staggered expression of the full set of seven-striped PRGs generating unique ‘double-segment’ codes to direct the formation of body segments (Gergen et al., 1986; Graham et al., 2019; Scott and Carroll, 1987). Many of the *Drosophila* PRGs transition to segmental expression as development proceeds but mutant phenotypes reveal the earliest roles of these genes in PR-patterning: roughly half-sized mutant embryos, missing alternate segments (Wieschaus and Nüsslein-Volhard, 2016).

**Figure 1.**
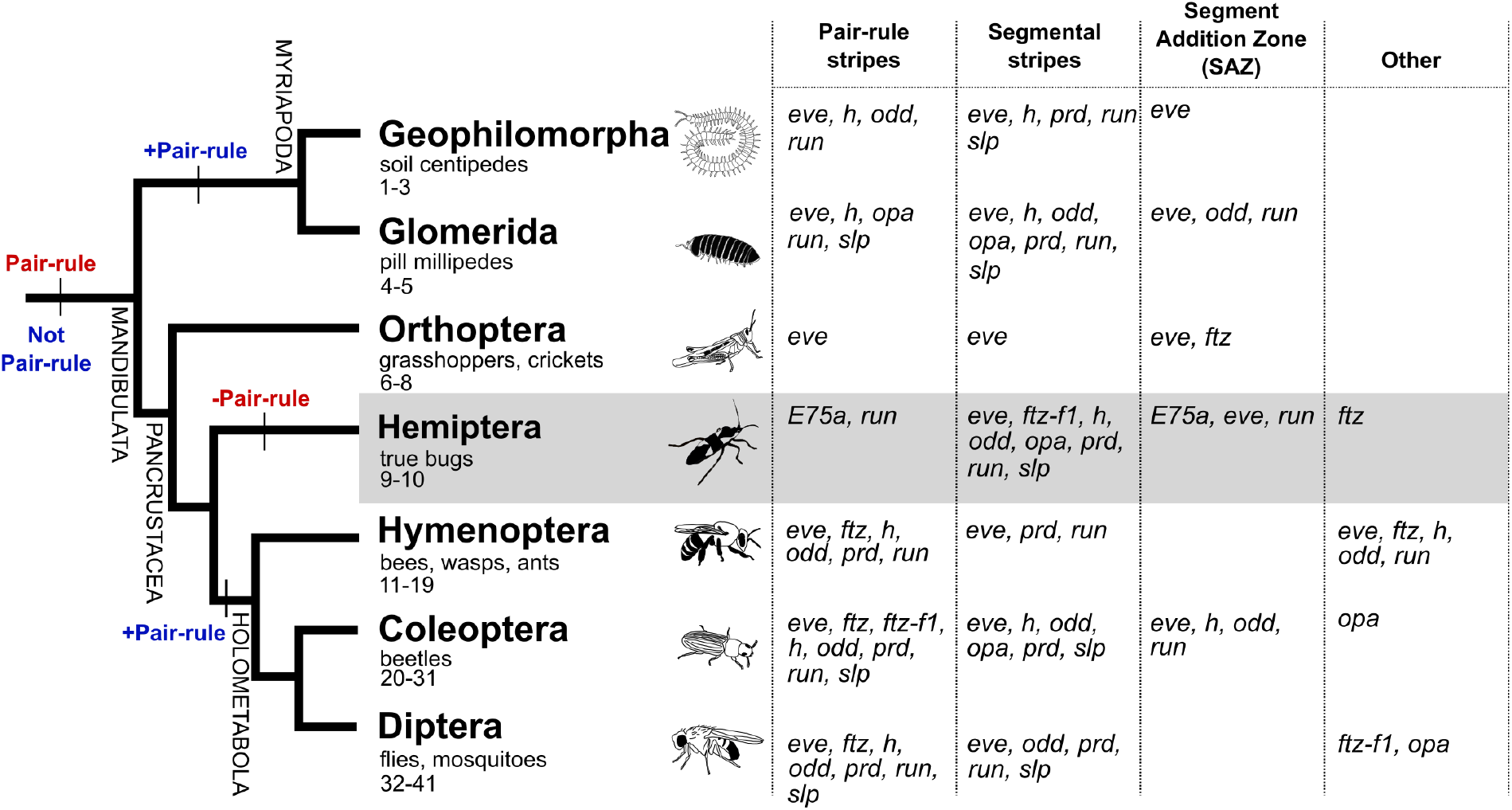
Models for the ancestral origin of PR-patterning. A simplified cladogram of arthropods is shown with segmentation-related expression patterns of PRG-orthologs in various insect and myriapod orders indicated as pair-rule-like, segmental (in every segment), SAZ (broad expression in the segment addition zone), or other. Non-segmentation-related expression patterns, such as expression in the nervous system, are not included. *Oncopeltus* is situated in the shaded region. Two hypotheses regarding the evolution of PR-like expression for the PRG-orthologs are shown in red and blue on the tree. Red) The ancestor of all arthropods exhibited PR-expression of the PRG-orthologs, which was subsequently lost in the lineage leading to *Oncopeltus*. Blue) PRG-orthologs were not expressed in a pair-rule manner in the arthropod ancestor, and PR-expression of these genes was gained independently in the lineages leading to myriapods and holometabolous insects. References for the expression patterns summarized here are detailed in Table S1.

Since all insects are segmented, the gene regulatory logic underlying segmentation might be wholly conserved. However, *Drosophila* are long-germ insects with all parasegments patterned more or less simultaneously at blastoderm. This mode of development is derived and only found among holometabolous insects, where it independently arose multiple times (Davis and Patel, 2002; Liu and Kaufman, 2005b). In contrast, most insect groups add segments sequentially after the blastoderm stage (‘sequential segmentation’), from the posterior end of the germband, a region known as the growth zone or segment addition zone (SAZ) (reviewed in (Davis and Patel, 2002; Liu and Kaufman, 2005b)). Thus, the *Drosophila*-like PR-patterning of a double-segment unit might be restricted to simultaneously segmenting species. However, PR-like expression patterns—defined as stripes of gene expression in the primordia of alternate (every-other) segmental units—have been observed in one or more sequentially segmenting species for orthologs of each of the nine *Drosophila* PRGs (PRG-orthologs): *ftz*, *fushi tarazu factor-1* (*ftz-f1*), *eve*, *odd skipped (odd), runt (run), hairy (h), odd paired (opa), paired (prd)* and *sloppy paired (slp)*. Within holometabolous insects, the expression and function of the complete set of orthologs of *Drosophila* PRGs has been examined in two species, the sequentially segmenting beetles *Tribolium castaneum* and *Dermestes maculatus* (Choe and Brown, 2007; Xiang et al., 2015; Xiang et al., 2017). These analyses, together with studies of selected PRG-orthologs in a handful of other holometabolous insects (Grbić and Strand, 1998; Kraft and Jäckle, 1994; Nakao, 2010; Nakao, 2015; Rosenberg et al., 2014), suggest that a role for some PRG-orthologs in *Drosophila*-style PR-patterning is shared among holometabolous insects, even those with sequential segmentation (Figure 1). However, other members of this gene set have changed in expression and/or function within Holometabola (Choe et al., 2017; Clark and Peel, 2018; Heffer et al., 2013a; Heffer et al., 2013b). For example, *ftz-f1* is expressed ubiquitously in *Drosophila* but in stripes in beetles (Heffer et al., 2013b; Xiang et al., 2017) and several PRG-orthologs are expressed in the segment addition zone (SAZ) in sequentially segmenting species as components of a vertebrate-like clock and wave mechanism, in addition to being expressed in PR-stripes (El-Sherif et al., 2012; Sarrazin et al., 2012).

In contrast to studies in holometabolous insects, there has been less focus on PRG expression or function in hemimetabolous insects. PR-like expression of *eve* was observed in a cricket (Mito et al., 2006; Mito et al., 2007) while in grasshoppers, *eve* and *ftz* have distinctly non-PR-like expression, both being expressed in the SAZ (Dawes et al., 1994; Patel et al., 1992). Also different from holometabolous species, *h* is expressed segmentally in a cockroach (Pueyo et al., 2008). Interestingly, PR-like expression of PRG-orthologs has been observed in evolutionarily distant, non-insect arthropods. For example, striped expression at half the frequency of segmental stripes (sometimes referred to as “double segment periodicity”), has been observed for several PRG-orthologs in a centipede (Chipman and Akam, 2008; Chipman et al., 2004; Green and Akam, 2013) and the expression of *prd* in spider mites is suggestive of modulation by a PR-like regulator (Dearden et al., 2002). These findings suggest two evolutionary hypotheses: PR-expression of PRGs arose independently in holometabolous insects and myriapods (Fig. 1, blue), or it was ancestral and lost in some hemimetabolous species (Fig. 1, red). If PR-like expression of PRGs was not ancestral (Fig. 1, blue), segmental expression of the PRG-orthologs may be the ancestral state. This scenario is supported by the observation that expression of five *Drosophila* PRGs evolves from a seven stripe PR-pattern to a fourteen-stripe, segmental pattern, either by stripe splitting or by de novo addition of a second set of seven stripes (Fig. 1). In either case, these results suggest extensive rewiring of segmentation networks in arthropods. Since it is clear that individual members of the PRG-set can vary in function without loss of PR-patterning per se (see above), it is necessary to examine the whole set of PRGs in diverse taxa before making broader conclusions about gain or loss of this patterning mechanism.

*Oncopeltus fasciatus* belongs to the order Hemiptera, a close outgroup to the Holometabola (Misof et al., 2014; Yeates et al., 2012). PR-like expression was observed in *Oncopeltus* embryos for the gene *E75A* and RNAi resulted in fusion of neighboring segments, demonstrating the existence of an underlying PR-like pre-patterning mechanism in this species (Erezyilmaz et al., 2009). However, *E75A* does not have PR-like expression or function in *Drosophila* (Bialecki et al., 2002; Buszczak et al., 1999; Segraves and Hogness, 1990). In *Oncopeltus, eve* is expressed in stripes in every segment (“segmental expression”) of the blastoderm and later in the SAZ (Liu and Kaufman, 2005a). Here we isolated and examined the expression of all nine orthologs of the *Drosophila* PRGs in *Oncopeltus (Of-*PRG orthologs), and seven paralogs of these genes, and compared their expression to the only known *Of*-PRG*, E75A.* Despite the fact that *Of*-PRG-orthologs are all expressed during the stages at which *Oncopeltus* specify segments, only one (*Of-run*) is expressed in a pattern reminiscent of *Drosophila* PRGs. Most others are expressed in segmentally reiterated patterns, either in nascent segments in the anterior SAZ or in mature segments of the germband. In keeping with this, PR-like defects were not seen after RNAi-mediated knockdown of *Of*-PRG-orthologs. These results suggest that, while PR-patterning *per se* is retained in *Oncopeltus*, extensive re-writing has occurred such that the genes responsible for this pre-pattern are different from those in *Drosophila*.

## Results

### *Isolation of* Oncopeltus *orthologs of* Drosophila *pair-rule genes*

Orthologs of the nine *Drosophila* pair-rule genes (PRGs), referred to throughout as *Of-*PRG orthologs, were isolated and gene structures determined (Fig. S1), combining experimental data with information from the *Oncopeltus* genome (Panfilio et al., 2019). Because the search criteria were designed to identify all potential orthologs, we identified multiple gene family members in most cases. All matches for each PRG were subjected to phylogenetic analysis to determine which, if any, were the ortholog of interest (Fig. S2). This analysis identified orthologs of *odd-skipped* family members *odd*, *sob* and *bowl*; one *opa* ortholog; *prd/Pax3/7* orthologs *prd* and *gooseberry*; four Runt domain family members, *run, lozenge, runxA*, and *runxB; h* family members*, h* and *deadpan;* one *slp* ortholog with 65% and 56% identity in the forkhead domain to *Dmel-*Slp1 and *Dmel-*Slp2; and a single copy of *ftz-f1*, with genomic sequence on four scaffolds that were merged after experimental verification.

For *Of-ftz*, three sequences encoding a homeodomain were isolated (Figs. S1, S3). *Of-ftz-A* (788 bp), *Of-ftz-B* (443 bp), and *Of-ftz-C* (965 bp). One of these, *Of-ftz-A*, was also found by RNA-seq (Ewen-Campen et al., 2011) and confirmed by us using 5’RACE. These sequences overlap in a region encoding a full-length, Ftz-family homeodomain. Upstream of the homeodomain, only one of these sequences, *Of-ftz-C*, appears to have a complete open reading frame; for this sequence, an HDWM appears to replace the YPWM motif seen in Hox proteins and homeotic-type Ftz proteins (Johnson et al., 1995). Further, the characteristic Ftz N-terminal arm (Heffer et al., 2010; Telford, 2000) differs from other Ftz proteins in arthropods. Of note is the substitution at position 4 of the homeodomain: *Of-ftz* encodes a lysine at this position, while all other arthropod Ftz examined share a serine or threonine (Heffer et al., 2010; Telford, 2000). The crystal structure of the *Drosophila* Engrailed homeodomain suggests that arginines at homeodomain N-terminal arm positions 3 and 5 make direct contact with DNA; these residues are conserved in *Of-*Ftz (Kissinger et al., 1990). For the shorter sequences (-*A* and *–B*), neither includes a start codon and stop codons are present in all three reading frames (Fig. S1 and S3). In addition, *Of-ftz-A* appears to include a 300 bp unprocessed intron; canonical GT-AG splice sites were found flanking an unaligned region of the sequence (Fig. S3). The sequences of all three *Of-ftz* homeoboxes match 100% at the nucleotide level and align to scaffold 2747 of the genome assembly. The portion of *Of-ftz-C* 5’ of the homeobox aligns to scaffold 1144. Despite the *Hox* complex being greatly fragmented in the current *Oncopeltus* genome assembly, a partial *Of-Scr* sequence was annotated on this scaffold. Future experiments will determine whether the three *Of-ftz* sequences isolated are different isoforms of one *Of-ftz* gene or products of distinct *Of-ftz* paralogs.

### Temporal expression of Of-PRG orthologs

Genes playing roles in PR-patterning are expected to be expressed first at the blastoderm stage and then throughout germband elongation, as segments are specified. Based on SYTOX green nuclear staining (Fig. S4), genes involved in segmentation should be expressed at 24-48 hours after egg laying (AEL), which includes late blastoderm through early germband elongation. RT-PCR spanning the first five days of *Oncopeltus* development was used to determine whether *Of*-PRG orthologs are expressed at the right time to be involved in segmentation (Fig. S5). For all experiments, *Of-actin* was simultaneously amplified as an internal positive control (Fig. S5). Due to its verified role as a PRG in *Oncopeltus* (Erezyilmaz et al., 2009), the expression profile of *Of-E75A*—a clear peak at 24-48h (AEL)—served as a guide for PR-like expression (Fig. S5A). Expression of *Of-eve* was detected at 0-24 h AEL, with highest levels at 24-48 h AEL and slightly lower levels after this (Fig. S5B). *Of-odd* and *Of-slp* expression were highest at 24-72 h AEL, from the blastoderm stage through germband extension (Fig. S5C,D), while *Of-slp* expression continued through the fifth day of embryonic development. *Of-run* expression was also highest at 24-48 h AEL, with attenuated expression for two more days (Fig. S5E). *Of-ftz-f1* showed fairly consistent expression for all time points, including at 0-24 h AEL (Fig. S5F), possibly reflecting maternal deposition, seen for *Drosophila ftz-f1* (Guichet et al., 1997; Yu et al., 1997). *Of-ftz* expression appears to be highest in 0-24 h AEL embryos and expression fades thereafter (Fig. S2G). Time courses for *Of-h, -prd*, and -*opa* expression were similar, with consistent expression detected 24-120 h AEL (Fig. S5H-J). Their continued expression after germband extension suggests additional roles later in development. In sum, all orthologs examined were detected at 24-48 h AEL, when highest expression of *Of-E75A* was also observed, consistent with roles in segmentation, although distinct profiles were seen for each gene.

### *Of-run* is expressed in a pair-rule-like manner during embryonic development

To determine whether *Of*-PRG-orthologs are expressed in PR-like spatial patterns, i.e., in stripes in the primordia of every-other body segment, whole-mount in situ hybridization was carried out. *Of-E75A*, expressed in PR-stripes (Erezyilmaz et al., 2009) serves as a positive control. *Of-E75A* was initially expressed in two stripes straddling the middle of the blastoderm-stage embryo, the anterior stripe being broad and diffuse and the posterior stripe much narrower (Fig. 2A). Slightly later, three stripes were observed; this third stripe was added from the posterior, at first diffuse and later narrowing (Fig. 2B, C). A fourth stripe was observed just before gastrulation (Fig. 2D). As gastrulation proceeded, all four blastoderm-stage stripes moved toward the posterior (Fig. 2E). In early germbands, a broad stripe was observed in the SAZ with a narrower stripe just anterior (Fig. 2F). In later germbands, a pair of stripes was seen in the anterior portion of the SAZ, with another weak stripe present in the segmental primordia (Fig. 2G).

**Figure 2.**
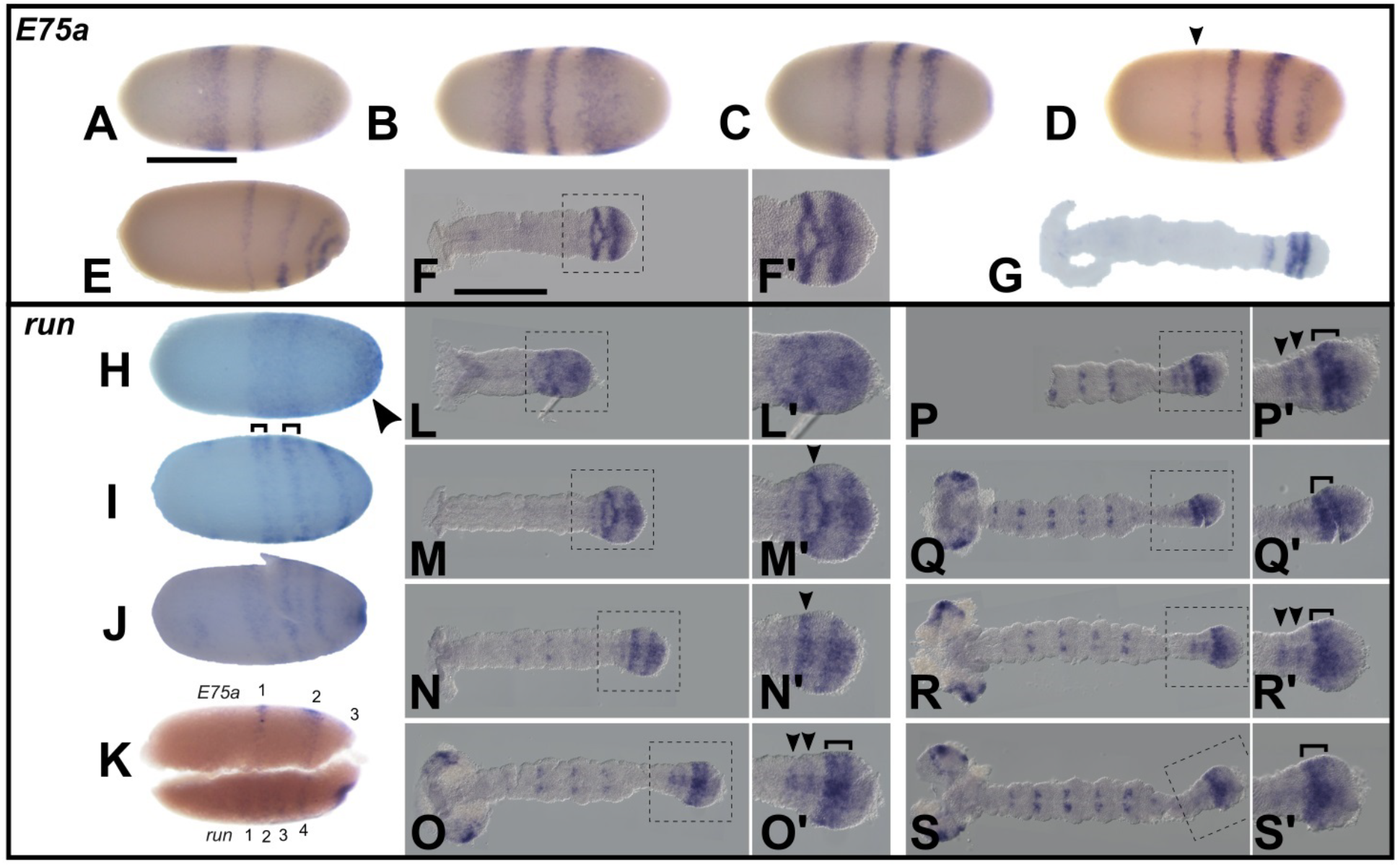
*Of-E75A* and *Of-run* are expressed in a pair-rule-like pattern. A-G) *E75A* expression, H-S) *run* expression. A-E) *E75A* was first observed as two stripes in the blastoderm. By germband invagination, four stripes were observed; F, F’) Early germbands show one stripe of expression anterior to the SAZ, and a broader stripe just posterior; G) A later germband with one faint stripe in segmented germband and a pair of stripe in the anterior SAZ; H) *run* was first observed in two centralized stripes and a posterior cap (arrowhead); I) The two anteriormost stripes appear to split (brackets) while the posterior cap resolves into a stripe; J) Five stripes observed at the onset of germband invagination; K) Three stripes of *E75A* visible in a blastoderm-stage embryo with four *run* stripes; it is possible that the posteriormost stripes have already invaginated at this point (*E75A* stripe 4, *run* stripe 5); L-S’) Dynamic SAZ expression was observed through anatrepsis in early germbands, including in some cases pairs of stripes (arrowheads) arising from the SAZ, where two-segment-wide stripes (brackets) were observed, similar to *E75A*. Embryos are oriented anterior left. Scale bars correspond to approximately 0.5 mm.

Of all the *Of*-PRG orthologs examined*, Of-run* expression was the most PR-like (Fig. 2H-S’). *Of-run* was first detected in blastoderm-stage embryos, in two diffuse stripes in the center of the embryo (Fig. 2H), as well as in a posterior ‘cap’ (arrowhead, Fig. 2H), similar to *run* expression in beetles (Choe et al., 2006; Xiang et al., 2017). At the beginning of gastrulation, stripes 1 and 2 split, while a third primary stripe resolved from the posterior cap generating five stripes prior to gastrulation (Fig. 2I, J). The presence of two thick stripes, each of which split, is reminiscent of PR-gene expression for *prd* in *Drosophila*, and *eve, h*, and *prd* in *Dermestes* (Kilchherr et al., 1986; Xiang et al., 2017). However, this pattern differs from *Of-E75A* which displays strict, alternate segment, PR-like expression, with stripes never splitting. To compare expression of these genes, embryos were bisected and halves were examined for either *Of-run* or *Of-E75A* expression. At late blastoderm, when two *Of*-*E75A* stripes were detectable, four *run* stripes were present (Fig. 2K).

During germband elongation, the *Of-run* posterior SAZ ‘cap’ persisted (Fig. 2L-N). A stripe just anterior to this SAZ expression was also observed (Fig. 2M’, N’ arrowhead). As abdominal segments were added, dynamic SAZ expression was evident (Fig. 2L’-S’). In some germbands, only one stripe was observed in the anterior SAZ, and *Of-run* expression in the posterior SAZ clearly extended to the posterior edge of the germband. In others, two stripes were evident in the anterior SAZ and a broad stripe, appearing to span the width of two segments, was observed just posterior (Fig. 2O, P, R), while expression in the germband’s posterior extreme had cleared. Expression was also seen in later germbands in head lobes, and in segmentally reiterated pairs of dots around the central midline in thoracic and abdominal segments (Fig. 2Q-S). In sum, *Of-run* appears to initiate expression in a PR-like register in blastoderm embryos, with stripes splitting to generate a segmentally reiterated pattern. *Of-run* is also expressed dynamically in the SAZ, similar to *run* expression in other sequentially-segmenting species. We classify *Of-run* expression as PR-like because of the initial broad stripes that split at blastoderm, as well as the appearance of stripes two-segments wide in the germband. However, a clear set of PR alternate-segment stripes, as seen for *Of-E75A*, was not observed.

### Of-prd *and* –odd *are expressed segmentally*

Segmental expression of *Of-eve* was reported by others but is included here for completeness (Liu and Kaufman, 2005a). *Of-eve* expression was first seen in a broad domain covering about 40-100% egg length (0%, anterior pole) (Fig. 3A). This domain then split into five stripes and a posterior cap, presumably by loss of transcripts from the inter-stripe regions (Fig. 3B). At the start of germband invagination, very weak stripes were observed, with a possible sixth stripe at the far posterior (Fig. 3C). In early germbands, tightly packed stripes in and around the SAZ were observed (Fig. 3D). Six blastoderm stripes are expected for genes expressed in the primorida of every segment, as is *Of-invected (Of-inv)*, a segmentally expressed gene. Note: *Of-inv* was originally thought to be *Of-engrailed (en)* (Genbank with accession number AY460340.1 (Liu and Kaufman, 2004); it has since been recognized as *invected* (Auman et al., 2017).

**Figure 3.**
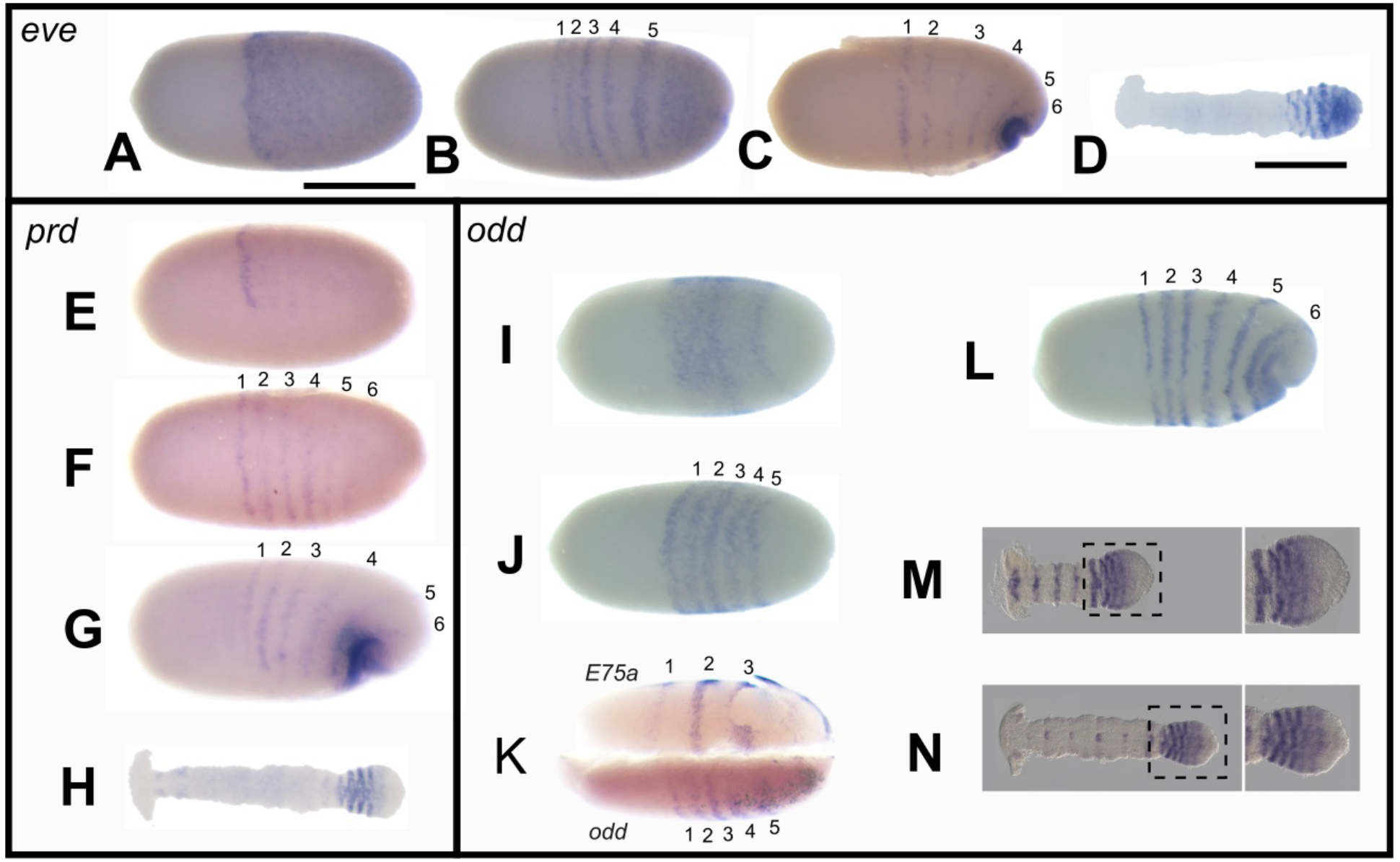
Of-*eve*, Of-*prd*, and Of-*odd* are expressed segmentally. A-D) *eve* expression, E-H) *prd* expression, I-N) *odd* expression. A-C) *eve* expression resolves into 6 stripes from an earlier broad domain of posterior expression in blastoderm-stage embryos; D) *eve* expression in the germband was limited to the SAZ, with a posterior cap and tightly packed segmental stripes in the anterior SAZ; E-G) *prd* stripe 1 arose first early blastoderm-stage embryos, followed by five additional stripes; H) 3-4 *prd* stripes were observed in the anterior SAZ in early germbands; I-J) Five *odd* stripes were observed in early blastoderm-stage embryos, resolving from an earlier broad domain; K) Three *E75A* stripes and five *odd* stripes were observed in a single bisected blastoderm-stage embryo (top half: *E75A*, bottom half: *odd*); L) a total of six *odd* stripes were seen at the onset of germband invagination; M, N) in germbands, *odd* was observed in stripes in the anterior SAZ; stripes in the mature segments were more apparent in earlier germbands. Embryos are oriented anterior left. Scale bars correspond to approximately 0.5 mm.

*Of-prd* was first detected at the blastoderm stage, in a dark narrow stripe at ~40% egg length (Fig. 3E). Two or three very light stripes just posterior to this were barely visible. This first stripe likely corresponds to the mandibular segment; the appearance of the mandibular *prd* stripe before others was observed in other insects (Choe and Brown, 2007; Osborne and Dearden, 2005; Xiang et al., 2015). At later blastoderm, six *Of-prd* stripes were observed; the posterior-most two stripes appeared much lighter than the presumably older, more anterior stripes (Fig. 3F). The stripes moved posteriorly as the germband invaginated (Fig. 3G). In early germbands, dots of expression were seen along the midline, indicating *prd* expression in central nervous system (CNS) development as in other arthropods (Davis et al., 2005; Osborne and Dearden, 2005). A group of four new tightly packed *Of-prd* stripes arose in the anterior SAZ similar to *Of-eve*, but with expression notably absent from the posterior SAZ (Fig. 3H). The register of these nascent stripes in the germband, along with the presence of six closely spaced stripes at blastoderm, demonstrate that *Of-prd* is expressed segmentally and not in a PR-like fashion.

*Of-odd* was first detected at blastoderm as a broad stripe at ~40-80% egg length, with 2-3 stripes resolving from this domain (Fig. 3I). Five narrow stripes appeared as the earlier diffuse expression cleared (Fig. 3J). Occasionally, a strong-weak alternation of these stripes was observed, possibly reflecting modulation of expression by a regulator expressed in a PR-pattern. An analogous alternate strong-weak pattern of stripes is seen for *Drosophila en*, likely reflecting regulation by different PRGs in alternate sets of stripes (DiNardo et al., 1985). To further test whether *Of-odd* stripes have segmental or PR-register, expression of *Of-odd* and *Of-E75A* were compared in bisected embryos. It is clear that there were two *Of-odd* stripes (bottom half of embryo) for every *Of-E75A* stripe (top half of same embryo) at this stage (Fig. 3K). In slightly older embryos after the germband started to invaginate, a sixth stripe arose at the posterior (Fig. 3L). In mature segments of the germband, expression in the presumptive mandibular through T2 segments gradually restricted to dots along the midline and one stripe in the newest mature segment (Fig. 3M). In the anterior SAZ, a group of four stripes, similar to *Of-prd*, was observed. In later germbands, a similar pattern was observed in the SAZ, but expression had faded in the segmented germband (Fig. 3N).

In sum, *Of-prd*, *-odd*, and *–eve* are expressed segmentally in blastoderm and germband stage embryos. For *Of-prd* and *Of-odd*, a cluster of stripes was observed in the anterior SAZ. The posterior cap of *odd* expression seen in beetles was not observed for *Of-odd* (Choe et al., 2006; Xiang et al., 2017). Overall, no hint of expression in a PR-like register was seen for *Of-prd*, *-odd*, or *–eve*.

### *Persistent segmental expression of* Of-opa *and* –slp

*Of-opa* expression was first detected in late blastoderm-stage embryos as six stripes on the lateral plates (Fig. 4A). In early germbands, before all six blastoderm stripes had invaginated, *Of-opa* stripes appeared in each segment (Fig. 4B). In later germbands, the six most anterior stripes had invaginated and persisted in each segment with weak expression just anterior to the SAZ (Fig. 4C). This persistent expression is different from *Of-odd* or -*prd*, where the 6 segmental stripes in the blastoderm eventually fade after those segment primordia become part of the germband.

**Figure 4.**
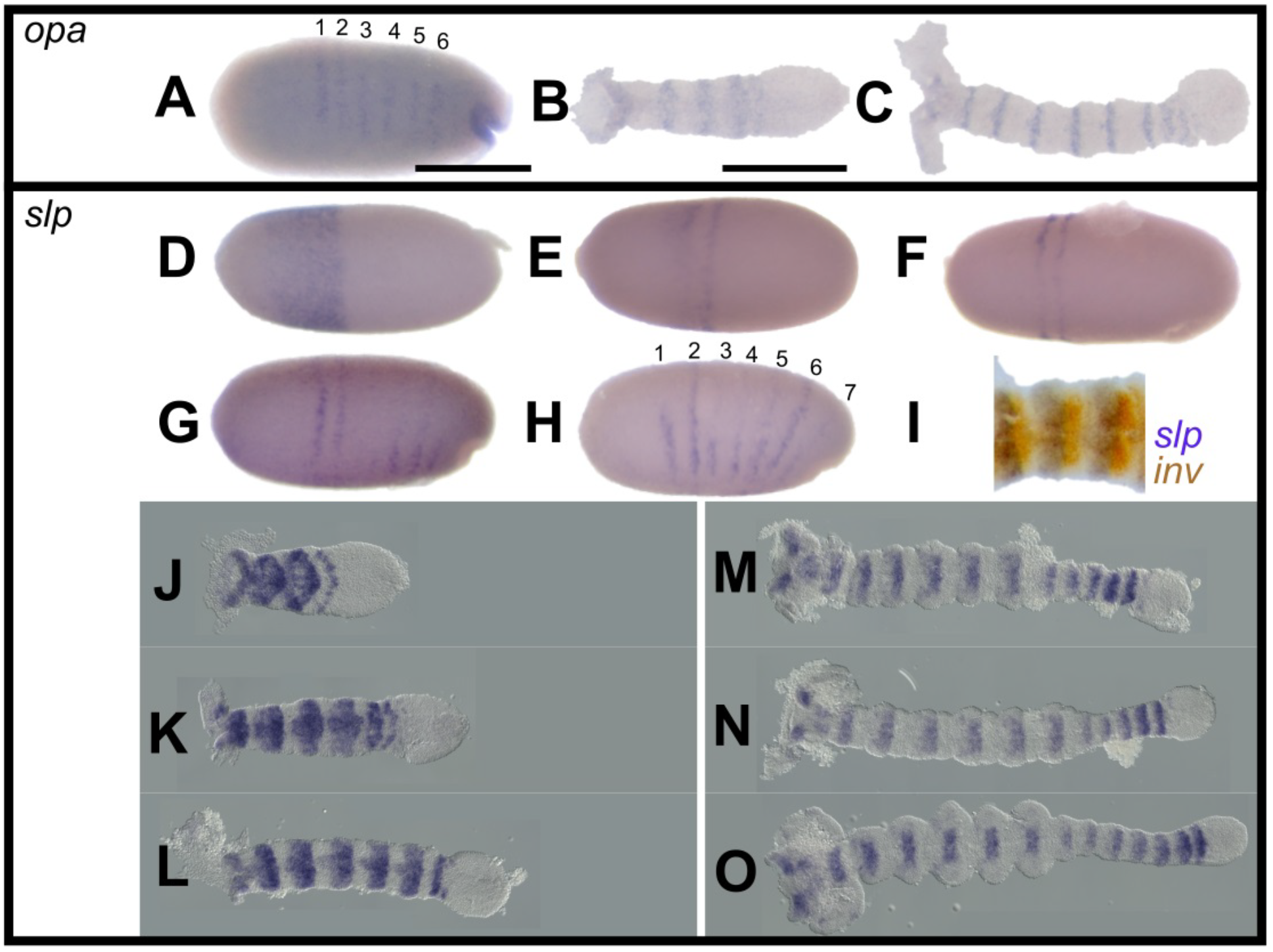
*Of-opa* and *Of-slp* are expressed in persistent segmental stripes. A-C) *opa* expression, D-O) *slp* expression. A) A total of six *opa* stripes were seen at blastoderm; B-C) *opa* stripes persisted in the mature segments of the early germband; D) early *slp* expression observed in the anterior half of an early blastoderm-stage embryo; E-F) the two anteriormost stripes arose first in the blastoderm; G-H) a total of seven *slp* stripes observed in the blastoderm; I) double stain with *slp* (purple) and *en* (orange) in an early germband; J-O) *slp* was seen in each mature segment of the germband during elongation, and was notably absent from the SAZ. Embryos are oriented anterior left. Scale bars correspond to approximately 0.5 mm.

*Of-slp* was first detected at blastoderm in a broad domain covering ~0-40% egg length, in a pattern complementary to *Of-eve* (Fig. 4D). In slightly later embryos, two stripes had emerged at the posterior boundary of this broad domain as the most anterior expression started to clear (Fig. 4E). This anterior expression had completely cleared by later blastoderm-stages, leaving two close stripes at ~30% egg length (Fig. 4F). A total of seven stripes were observed at blastoderm (Fig. 4G, H). In early germbands, double staining revealed *Of-slp* expression spanning the mediolateral width of each segment, anterior to each *inv* stripe (Fig. 4I). As germband elongation continued, a broad stripe of *Of-slp* expression persisted in each mature segment (Fig. 4J-O). These segmental stripes have clearly defined posterior borders but more diffuse anterior boundaries.

In sum, both *Of-opa* and *Of-slp* are expressed in segmental stripes in the blastoderm which persist through germband elongation. These genes differ from *Of-eve* –*odd*, and-*prd* in that *Of-opa* and *–slp* were not detected in the SAZ and their striped expression persisted in the elongated germband, similar to *slp* in *D. maculatus* (Xiang et al., 2017).

### Of-h, -ftz-f1, *and* -ftz *have unique features*

*Of-h* expression was first detected later than other PRG-orthologs, at late blastoderm-stage, in three faint stripes (Fig. 5A). These stripes were never observed to split, and this pattern was observed at much later stages than the three-stripe pattern of *Of-E75A* (note the invagination pore in Fig. 5A). These stripes are likely the earliest manifestation of the germband expression seen slightly later. In germbands, *Of-h* stripes were observed posterior to each *inv* stripe (Fig. 5B). Notably, expression of *Of-h* appeared to lag behind that of the other genes examined, as it was observed in older, more mature segments, rather than in the younger segments being generated from the SAZ, suggesting coordinate activation of *Of-h* stripes by gene(s) already expressed segmentally (Fig. 5C-E). Auman and Chipman (2018) also found *h* expression in two stripes in the anterior SAZ. In later germbands, new stripes were seen in the abdominal segments, in addition to expression in antennal segments (Fig. 5F). In fully extended germbands, expression in the abdomen faded away, and new expression was seen in labial through T3 segments (Fig. 5G). Later, two or three dots in thoracic appendage primordia were observed (Fig. 5H).

**Figure 5.**
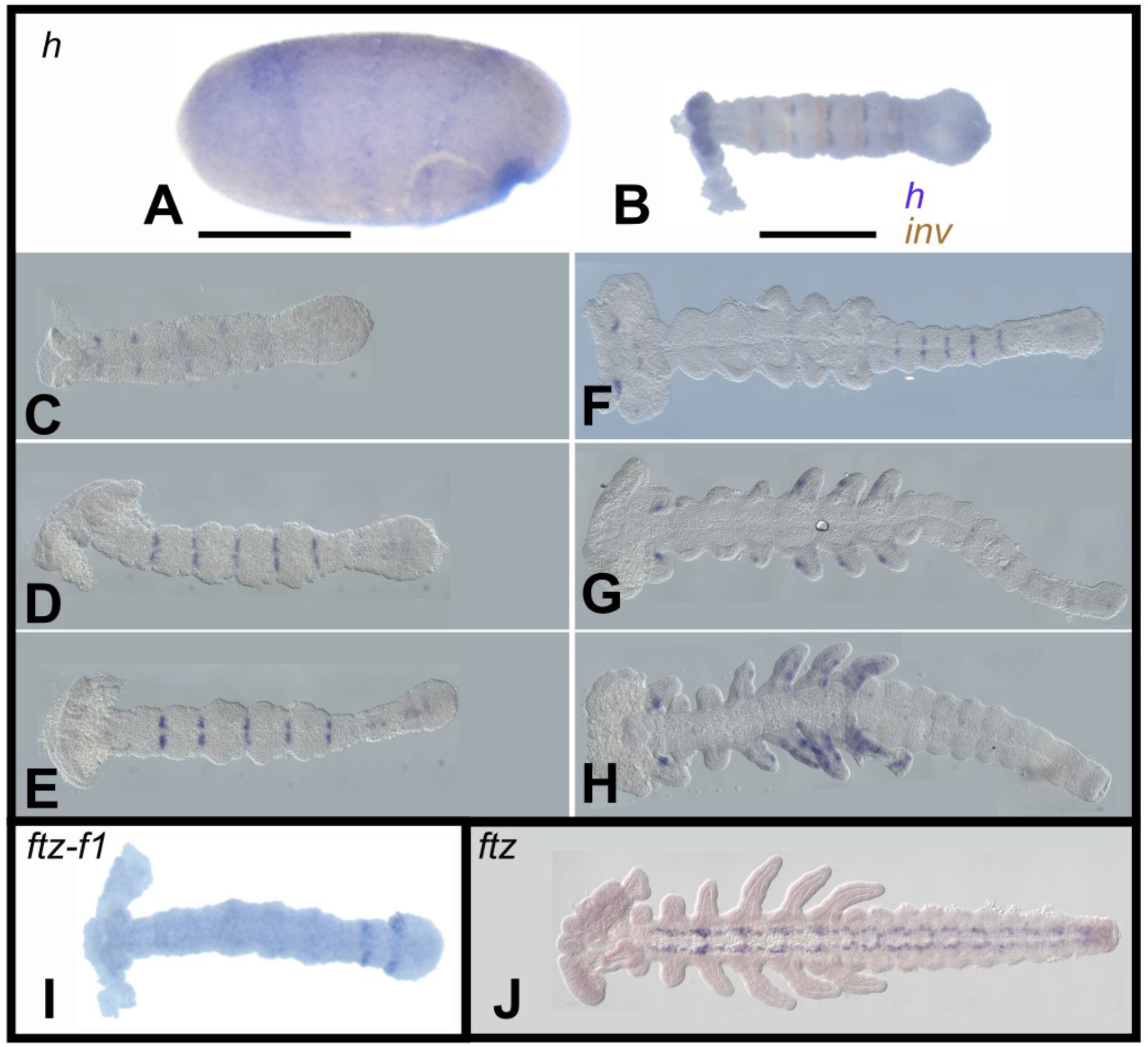
*Of-h* and *Of-ftz-f1* are expressed segmentally while *Of-ftz* shows conserved CNS expression. A-H) *h* expression, I) *ftz-f1* expression, J) *ftz* expression; A) At late blastoderm, *h* was observed in three stripes; B) double stain with *en* (orange) and *h* (purple); C-F) segmental stripes of *Of-h* were observed through germband elongation; G, H) in later germbands, *h* expression was observed in the limb primordia; I) *ftz-f1* was observed in 2 stripes just anterior to the SAZ in the elongating germband; J) *ftz* was observed along the midline in the presumptive CNS primordium in fully extended germbands. Embryos are oriented anterior left. Scale bars correspond to approximately 0.5 mm.

No blastoderm expression of *Of-ftz-f1* was detected. In germbands, two stripes were observed anterior to the SAZ during early abdominal segment addition; similar to *h*, these stripes were weaker along the central midline (Fig. 5I). Later, only one *ftz-f1* stripe near the SAZ was observed (data not shown). Double in situ hybridization with *Of*-*inv* revealed that *Of*-*ftz-f1 and Of-inv* segmental stripes are out of register with each other (data not shown).

No localized expression pattern was observed for *Of-ftz* during segmentation, using an *Of-ftz-C*-specific probe or a probe that would detect all three isoforms (Fig. S1). At later stages, segmental expression of *Of-ftz-C* was observed in groups of internal cells, presumably CNS, as has been seen for *ftz* in other species (Dawes et al., 1994; Heffer et al., 2013a) (Fig. 5J).

### Paralogs of *Of-*PRGs are not expressed in pair-rule-like patterns

Since functional divergence following gene duplication can lead to subfunctionalization of ancestral protein functions, it is possible that for any of the *Of-*PRGs, the ancestral gene encoded a protein with pair-rule function, and that this function was relegated to a different paralog in the lineage leading to *Oncopeltus* than that leading to *Drosophila.* To investigate whether *Of-*PRG paralogs are expressed in PR-like patterns, and thus retain potential to perform PR-functions, the timing of expression of paralogs was determined by RT-PCR. *gooseberry-neuro (gsb-n)* was not found in the *O. fasciatus* genome, nor in the *H. halys* genome (Fig. S2C), suggesting loss of this gene in Pentatomomorpha. All other paralogs investigated were located in the genome and gene identity was determined by phylogenetic analysis (Fig. S2). *runxB* could not be amplified using mixed stage 0-120 h AEL cDNA, nor could it be identified in the *Oncopeltus* embryonic transcriptome (Ewen-Campen et al., 2011); *sister of odd and bowl (sob), brother of odd with entrails limited (bowl), gooseberry (gsb), runxA, lozenge (lz)*, and *deadpan (dpn)* were all found to be expressed within this time frame. As segmentation occurs during the first three days of embryogenesis, *runxA*, which was found to be expressed 72-120 h AEL, was excluded from further analysis.

Expression patterns of *sob, bowl, lz, dpn*, and *gsb* were determined by in situ hybridization on 24-72 h AEL embryos. No patterned expression was observed for *lz* and *bowl*. *odd* paralog *Of-sob* was observed in an *Of*-*odd*-like pattern at blastoderm in six segmental stripes and later in stripes in the anterior SAZ (Fig. S6A-D). This *Of-sob* expression pattern was also found by Auman and Chipman (2018), who showed by comparison with *Of-eve* that *Of-sob* and *Of-odd* stripes overlap. Fully elongated germbands showed *Of*-*sob* expression in stripes in the appendages, suggesting a role in appendage patterning, as was shown for *sob* in *Tribolium* (Angelini et al., 2012). Expression of *Of-dpn, Of-h* paralog, was observed in the head lobes and later along the midline, possibly in CNS primordia (Fig.S6F-G). *Of-gsb, prd* paralog, was observed in one stripe at blastoderm similar to early *Of-prd* expression (compare Fig. S6H and Fig. 3E), and later in each mature segment of the elongating germband. Thus, none of the paralogs are expressed in PR-like patterns.

### Parental RNAi of some *Of-*PRGs results in severe disruption of *inv* expression

Parental RNAi (pRNAi) was performed to assess the function of each *Of-*PRG. qPCR on RNA extracted from pRNAi embryos was used to determine relative expression and verified knockdown of each gene targeted (Fig. S7). Expression of *Of-inv* in elongating germbands was used to assay segmentation phenotypes as loss of alternate *inv* stripes would be expected after loss of PR-function. RNAi knockdown of *odd* or *prd* resulted in severe defects (Fig. 6B, C). No thoracic or abdominal segmentation was apparent in these shortened embryos, suggesting overall loss of segmentation, more similar to what is seen for *Drosophila* segment polarity mutants than *Drosophila* pair-rule mutants. Consistent with this, in different RNAi embryos, *inv* was undetectable, detectable in partial stripes at only very low levels (Fig. 6B), or unpatterned throughout the embryo (Fig. 6C). These effects were seen throughout the embryo, without PR-like register, indicating that *Of-odd* and *Of-prd* impact *inv* stripes in all segments. While the *Of*-*prd* knockdown phenotype was fairly consistent (22/22 embryos, Fig. 6C, Fig. S8B), slightly more variation in phenotype was observed in *Of-odd*^pRNAi^ offspring. These differences appeared to broadly stratify with different replicates, indicating that changes in dsRNA integrity or amount of dsRNA injected may cause this variation. In some *odd*^*p*RNAi^ embryos, every *inv* stripe was nearly lost in every segment (15/29 embryos, Fig. 6B, Fig. S8A7-16) while in others, *inv* expression was less severely affected (14/29 embryos, Fig. S8A1-6). Auman and Chipman (2018) found fusion of maxillary and labial segments, as well as first and second thoracic segments after *odd* RNAi, which we did not observe. Unhatched *odd*^*p*RNAi^ offspring showed severe segmentation defects, often with all thoracic appendages fused (Fig. 6H), while unhatched *prd*^pRNAi^ offspring developed only heads (Fig. 6I). In contrast to *Of*-*odd* and *Of*-*prd* knockdown, striped *inv* expression was largely maintained in *Of*-*slp*^*p*RNAi^ embryos, although five rather than six stripes were consistently observed in the gnathal/thoracic region (23/23 embryos, Fig. 6D and Fig. S8C). Stripes appeared expanded, especially along the midline. Since *Of*-*slp* stripes are offset from *inv* stripes (Fig. 4I), this expansion suggests that *Of*-*slp* represses *inv* but without PR-like register. These embryos showed no clear morphological segmentation and failed to develop appendages (Fig. 6J), as found by Auman and Chipman (2018).

**Figure 6.**
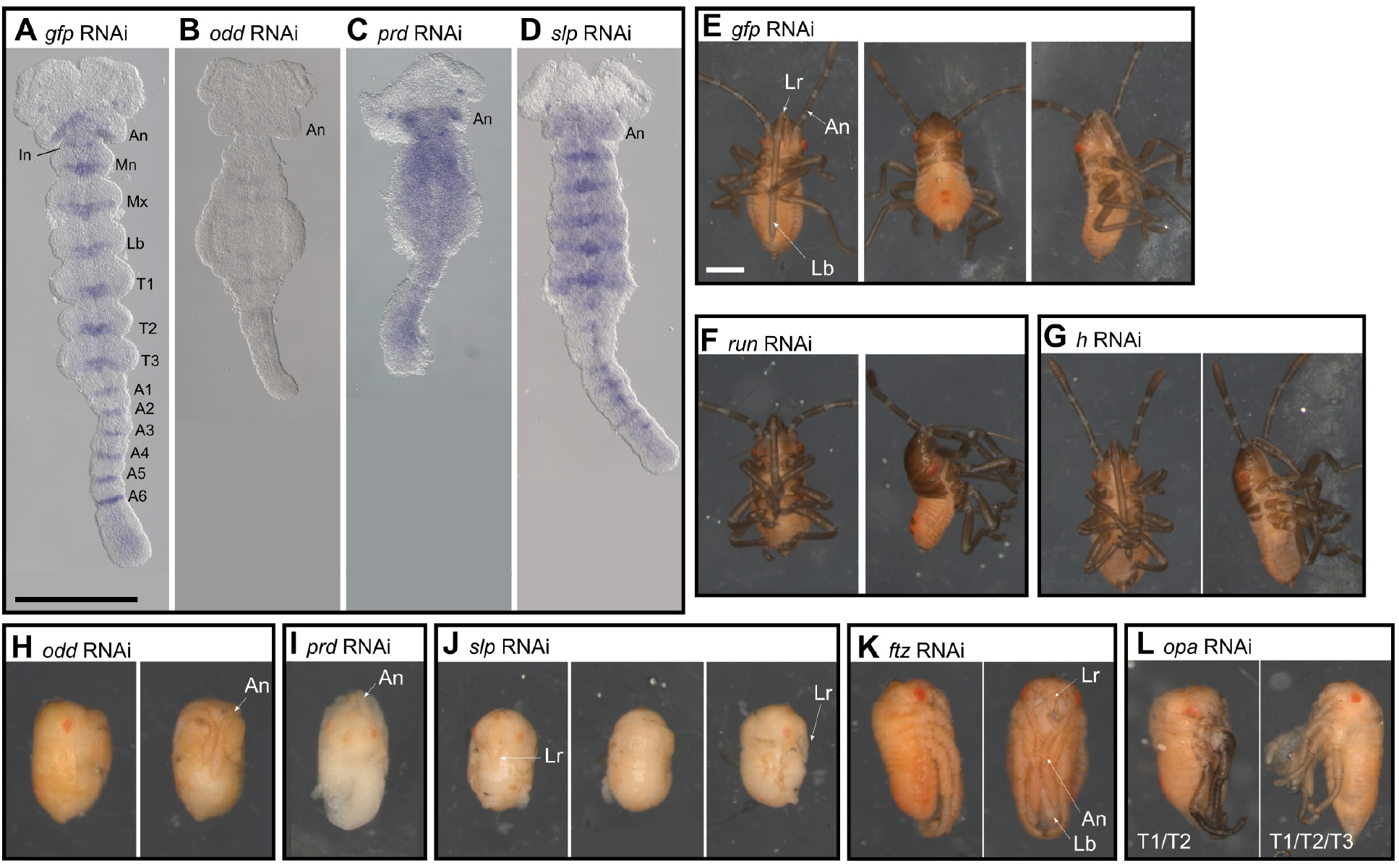
RNAi knockdown of *Of*-PRGs. Parental RNAi was performed. A) *gfp*^pRNAi^ was used as a negative control; B) *odd*^pRNAi^ resulted in near-total loss of each *inv* stripe; C) *prd*^pRNAi^ generated background levels of *inv*, detected throughout the embryo; D) *slp*^pRNAi^ disrupted *inv* stripe boundaries, especially along the midline, and suppressed appendage formation; E-H) *gfp, run*, and *h* pRNAi hatchlings appear wild-type-like; H) *odd*^pRNAi^ offspring show normal head and antennae, but lacks differentiated thoracic appendages and wild-type abdominal segmentation; I) truncated *prd*^pRNAi^ offspring with clearly developed head, but lacking remaining body segments (‘head-only’); J) *slp*^pRNAi^ offspring with distinct labrum, but lacking all gnathal and thoracic appendages and clearly defined abdominal segments; K) *ftz*^pRNAi^ offspring fail to hatch due to malformed gnathal appendages; L) *opa*^pRNAi^ offspring display irregular thoracic appendage fusions. *An*: antennal segment or antenna; *In:* intercalary segment; *Mn:* mandibular segment; *Mx:* maxillary segment; *Lb:* labial segment or labium; *Lr:* labrum. Scale bars correspond to approximately 0.5 mm.

Females injected with dsRNA targeting *Of*-*ftz-f1* failed to lay eggs, and dissection revealed that injected females’ ovaries were abnormal, containing oocytes of about the same size along the length of the ovariole (Fig. S9), similar to what was seen after *Tribolium* and *Dermestes ftz-f1*^RNAi^ (Heffer et al., 2013b; Xiang et al., 2017; Xu et al., 2010). RNAi knockdown of *Of*-PRGs *ftz, h, run*, and *opa* did not impact *Of-inv* expression compared to *gfp* controls (47/47, 8/8, 27/27, 166/167, and 37/37 embryos examined, respectively). While hatch rates were severely suppressed for all concentrations of *Of*-*opa* dsRNA tested, no effect was observed on *inv* stripes (Fig. S10). Examination of unhatched first instar offspring revealed irregular fusions of thoracic appendages (Fig. 6L), suggesting that *Of*-*opa* is required to maintain segment boundaries later in embryogenesis. For *Of*-*h, ftz* and *run*, hatch rates were only slightly lower than controls (Fig. S10). Cuticular defects were observed after *Of*-*ftz*^*p*RNAi^ (Fig. 6K) in which the maxillary and mandibular segments appeared undifferentiated, similar to *Of-Scr*^*pRNAi*^, suggesting possible off-target effects; however the labium appeared wild-type-like in contrast to the distinct labial to antenna or leg transformation observed after *Of*-*Scr* knockdown (Hughes and Kaufman, 2000). No defects in body patterning were observed in *Of*-*run* or -*h* pRNAi offspring (Fig. 6F-G). In most cases, more than one dsRNA sequence was tested at multiple concentrations for each gene. Levels of knockdown as measured by qPCR for these genes were similar to those for genes showing abnormal *inv* expression (*Of*-*odd, -prd*, and -*slp*), suggesting that RNAi was effective (Fig. S7). However, functional inference from RNAi is suggestive but not definitive; future experiments will be required to generate genomic deletions of these genes to more conclusively assess their function.

In sum, for those PRG-orthologs showing RNAi-defects in early embryos, all segments were affected equally. Thus, these genes do not appear to function in a PR-like manner in *Oncopeltus*.

## Discussion

We isolated and examined the expression patterns and functions of orthologs of the nine *Drosophila* PRGs, as well as their paralogs, in *Oncopeltus*. This is, to our knowledge, the first examination of the complete set of PRG-orthologs in a hemimetabolous insect. Using this candidate gene approach, we found that only one ortholog, *Of-run*, is expressed in a striped pattern reminiscent of *Drosophila* PRGs. However, unlike *Of-E75A*, which shows PR-like expression*, Of-run* was not observed in a striped pattern in alternate segments in the blastoderm, but rather in two broad stripes that then split, and no pair-rule defects were observed after knockdown. Most *Of-*PRG orthologs were expressed segmentally during blastoderm and germband development. The finding that all *inv* stripes were impacted after *Of-odd, -prd* or *-slp* RNAi knockdowns suggests that these genes are involved in the specification or boundary formation of every segment. This function resembles that of *Drosophila* segment polarity genes. Among the *Of*-PRG-orthologs expressed segmentally, distinct temporal and spatial classes were observed. *Of-eve, -odd*, and *–prd* were expressed in six segmental stripes in the blastoderm, and in germbands were expressed with segmental register in a transitional zone between the posterior SAZ and the segmented germband. This is of interest because segmental stripes appear to emanate directly from the SAZ. In contrast, *Tribolium* orthologs expressed in the SAZ emerge with PR-periodicity. *Of-opa* and -*slp* were observed in a similar segmental pattern in the blastoderm and were later seen as persistent stripes in each mature segment of the germband, but were absent from the SAZ. *Of-h* and *Of-ftz-f1* were observed in stripes only at later stages in the mature germband; these stripes appear to be more spatially restricted. *Of-ftz-f1* was never seen in more than two stripes anterior to the SAZ; *h* was present in sets of stripes, either in anterior or in posterior segments, just posterior to *Of-inv* in germbands. The temporal differences in expression of these genes, along with the increasingly restricted spatial localization of the stripes, suggests a regulatory hierarchy among them. If indeed PR-expression for this gene set was ancestral (Fig. 1, red), a coordinated shift in their expression patterns from PR-like to segmental could be explained by cross-regulatory interactions.

While this work was in progress, Auman and Chipman (2018) examined expression of *Of-odd, -opa, -slp, -h*, and *-run* during germband elongation and tested RNAi knockdown of *Of-odd* and *Of-slp*. Their results were largely similar to those shown here. Our study differs from their study in that we analyzed expression and function for the full set of *Of*-PRGs, including *Of-ftz*, *ftz-f1*, *prd*, as well as paralogs. Further, a major goal of our approach was to determine whether these genes were expressed in PR-like register. For this, we focused on examination of expression of each gene at the blastoderm stage, where stripe register can be determined clearly by comparison to the one known *Oncopeltus* gene expressed in a PR-like pattern, *Of-E75A* (Fig. 2). Comparison to *Of*-*E75A* expression, including examination of expression in bisected embryos (Fig. 2K, 3K) allowed us to make strong conclusions about the mode of expression of *Of*-PRG orthologs.

Are orthologs of *Drosophila* PRGs involved in an ancestral PR-mechanism? Orthologs of *Dmel*-PRGs are expressed in PR-like patterns in a number of holometabolous insects (Aranda et al., 2008; Choe et al., 2006; Rosenberg et al., 2014; Wilson and Dearden, 2012; Xiang et al., 2017), although expression of individual orthologs in different species varies (Fig 1), as mentioned in the Introduction. This variation among PRG-orthologs in Holometabola may reflect an evolutionary reshuffling of functions within one level of a regulatory hierarchy. In hemimetabolous insects the situation appears to be different: here we have found absence of PR-like expression or function for most of the PRG-orthologs and their paralogs. Though the full set of PRG-orthologs has not been examined in other hemimetabolous insects, there is variability in the degree of conservation of expression and function in those orthologs that have been studied. For example, in grasshoppers, neither *eve* nor *ftz* is expressed in stripes (Dawes et al., 1994; Patel et al., 1992), however *eve* is expressed in broad, PR-like stripes that split to generate segmental expression in the cricket (Fig. 1, (Mito et al., 2007). In contrast, an ancestral role for PRG orthologs in patterning a double segment repeat was suggested from studies of myriapods. In particular, in the centipede *Strigamia maritima*, orthologs of *odd (Sm-odr1)*, *eve, run*, and *h* are expressed in stripes that appear in alternate segmental intervals (Chipman and Akam, 2008; Chipman et al., 2004). There is also some evidence for PR-like expression of PRG-orthologs in the millipede *Glomeris marginata*, based on the finding that initial expression of a number of these genes (*eve*, *run*, *h*, *slp*, *opa*) is in double- or multiple-segment wide domains, which then split into segmental stripes (Janssen et al., 2012). We note that this differs from *Drosophila*, where no single PRG is expressed in a stripe spanning a double segment primordia; the widest PR-stripes are those of *prd*, which are broader than other PRGs but do not span an entire double segmental unit (Gutjahr et al., 1993). The “double segment” prepattern inferred for *Drosophila* PR-patterning arises from the complementary expression of different PRGs. Thus, these wide PRG-ortholog stripes are not necessarily reflective of a *Drosophila*-like mechanism. However, an underlying PR-mechanism is suggested by the fact that, as seen in the spider mite (Dearden et al., 2002), the *Glomeris prd*-ortholog is expressed in segmental stripes that are stronger in alternate segments (Janssen et al., 2012).

These findings highlight the distinction between PR-pre-patterning as a developmental mechanism, and the specific genes identified in *Drosophila* that control this process. PR-like pre-patterning occurs in *Oncopeltus* largely or wholly independently of the set of PRGs that play this role in holometabolous insects. *E75A*, which has no PR-like expression in *Drosophila* (Wilk et al., 2013), is expressed in the primoridia of alternate body segments in *Oncopeltus* ((Erezyilmaz et al., 2009), and Fig. 2), demonstrating that an underlying PR-type mechanism exists in *Oncopeltus.* Interestingly, PR-like expression in *Oncopeltus* was also seen for *Toll*-like genes, which are downstream targets of PRGs in *Drosophila* ((Benton et al., 2016; Graham et al., 2019; Paré et al., 2014) and data not shown). Since the *Drosophila* regulators of these *Toll* gene PR-patterns are non-PR in *Oncopeltus*, the regulatory interactions controlling their expression must have been re-wired in one of these lineages. Thus, while there is little overlap between *Oncopeltus* and *Drosophila* in the sets of genes responsible for the initial subdivision of the embryo into double segment repeats, PR-pre-patterning is part of the ‘regulatory logic’ by which the embryo is sequentially subdivided into increasingly specified units. The endpoint of this process—segment formation—is highly stable during evolution, but the regulatory pathways directing it have diverged, a phenomenon termed developmental systems drift (Haag, 2014; True and Haag, 2001), which we refer to as ‘phenotypic stability.’

We close with an historical note of interest. Before geneticist Peter Lawrence shifted the focus of his career to the *Drosophila* model system, he did extensive research on *Oncopeltus* segmentation (Lawrence, 1966; Lawrence, 1973; Lawrence and Green, 1975). Through clonal analysis of irradiated embryos, he showed that patterning of segments begins at the blastoderm stage of *Oncopeltus*. Given the complexity of PR-patterning in *Drosophila* and other holometabolous insects, we expect that, in addition to *E75A*, other genes with PR function remain to be discovered in *Oncopeltus*. It is therefore not a stretch to speculate that if Lawrence had continued his studies of segmentation in *Oncopeltus*, we might be referring to a whole different set of regulatory genes when thinking about PR-patterning in insects. This example is one of many that underscores the importance of studying diverse experimental systems to understand biodiversity at the regulatory level; limiting our studies to a handful of organisms inevitably biases our understanding of developmental processes.

## Materials and Methods

### Insect rearing

*Oncopeltus*, originally purchased from Carolina Biological and maintained in our lab for several years, were reared on a diet of water and sunflower seeds at 25±1°C, 50% RH, with a photoperiod of 16L:8D.

### Embryo in situ hybridization

Embryo fixation and in situ hybridization was carried out as described by Liu and Kaufman (2004) with modifications made by Ben-David and Chipman (2010) and by our lab (see SI). Antisense RNA probes were synthesized using digoxigenin or fluorescein RNA labeling mix (Roche) using standard methods. BCIP/NBT was used most often as the chromogenic substrate for the alkaline phosphatase; in the case of double in situs, BCIP/INT was used to produce an orange product. Since our original probes were designed to hybridize to conserved domains, we designed additional probes for *odd, prd, run*, and *slp* that were gene-specific to ensure that our results were not obscured by cross-hybridization with related genes. Gene-specificity was confirmed by conducting a BLAT search of the *Oncopeltus* genome using the probe sequence as the query; if only the sequence of the target gene matched the probe sequence, we considered the probe gene-specific. We confirmed with *odd, prd, run*, and *slp* that the gene-specific probes (Fig. S1, labeled probe 2 or 3) gave the same results as the less specific probes (Fig. S1, labeled probe 1).

### Parental RNAi

Double-stranded RNA for RNAi was synthesized using the Megascript T7 Transcription Kit (Invitrogen); dsRNA target regions are shown in blue in Fig. S1. Template for the RNA synthesis reaction was produced by PCR of cloned sequence fragments, using primers with the T7 promoter sequence at the 5’ ends. The RNA synthesis was allowed to proceed for 16 h at 37°C, then treated with TURBO DNase, and annealed in a thermocycler. Each 20 uL reaction of double-stranded RNA was precipitated with 30 uL of 7.5M lithium chloride and 250 uL cold ethanol and then resuspended in injection buffer (5 mM KCl, 0.1 mM phosphate buffer pH 6.8). A 1:100 dilution of each dsRNA was run on an agarose gel to check that dsRNA was indeed present and the expected size.

Anticipating possible variability in effect between different dsRNAs, we tested three different concentrations per dsRNA (10, 15, and 20 ug), injecting 2-3 females with each concentration. Embryos were collected daily and the hatch rate was tracked for 3-4 weeks post-injection, or until injected females died (Fig. S10). One of the longest living females, injected with 15 ug *prd* dsRNA, continued to lay clutches that failed to hatch 31 days after injection, a much longer penetrance than what has been reported previously in this species (Liu and Kaufman, 2004). The remaining experiments were performed with the lowest concentration of dsRNA that produced a noticeable effect on the hatch rate. For some dsRNAs (*h, run, opa*, and *ftz*) no effect on hatch rate was observed even at the highest concentration tested. For these, additional experiments were performed with the highest concentration. Adult females were injected with dsRNA corresponding to each *Of*-PRG-ortholog about a week after molting from L4 to adult. Injections were done in triplicate, with 3-5 females per group for each gene. Double-stranded *gfp* RNA was injected as a control. One day after injection, an equal number of males was added to each cage. Embryos were collected daily and divided such that some were allowed to hatch, some were fixed at 48-72 h AEL for subsequent staining by in situ hybridization with an *inv* probe, and some were frozen in TRIzol at −80°C at 24-48 h AEL (*odd, prd, slp, h, run, ftz* and *gfp*) or 35-50 h AEL (*opa* and *gfp*). A tighter staging of *opa* RNAi embryos was necessary to ensure embryos used for RNA extraction were expressing *opa.* An additional dsRNA from a different part of the gene sequence was tested for *prd, odd, slp, h, run*, and *opa* in a separate round of experiments (Fig. S1, red). Results were largely similar; defects seen for *slp* with this dsRNA were weaker; however, no defects were seen for *Of-odd* with this second dsRNA, which matched the *odd* 3’UTR. Thus it is possible that defects shown above and by Auman and Chipman (2018) reflect knockdown of both *Of-odd* and *–sob*, which appear to be expressed in the same pattern (compare Fig. 3I-N to Fig. S6A-D).

### Quantitative RT-PCR

qPCR was performed using 24-48 h AEL (*prd, odd, slp, ftz, h*, and *run*) or 35-50 h AEL (*opa*) RNA from RNAi embryos. RNA was extracted using TRIzol (Invitrogen), DNase-treated using the TURBO DNA-free kit (Invitrogen), and cDNA was transcribed using the iScript cDNA Synthesis kit (Bio-Rad). PCR was performed in a Roche LightCycler 480 real-time PCR machine, using the Luna Universal qPCR master mix (NEB). *TATA-box binding protein (Tbp*, found on scaffold 2359 of the *Oncopeltus* genome) was used as the reference gene as it was found to have the most stable expression of three candidate reference genes tested, and has been used as a reference gene in many other studies (Liang et al., 2014; Niu et al., 2014; Zhai et al., 2014). The 2^−ΔΔCT^ method was used to calculate fold change of gene expression relative to the *gfp* control. Statistical significance was determined by performing a Welch two-sample t-test in R with α = 0.05. All primer sequences are listed in Table S2.

### Phylogenetic analyses

To conduct phylogenetic analyses of our candidate ortholog sequences, alignments of orthologous sequences were generated using MUSCLE (Edgar, 2004). Alignments were then trimmed using Aliview (Larsson, 2014). Finally, phylogenetic analysis was carried out in TOPALi v2.5 using the Bayesian algorithm MrBayes (Milne et al., 2009; Ronquist and Huelsenbeck, 2003). Additional formatting of trees was performed in MEGA6 (Tamura et al., 2013).

## Acknowledgements

This work was supported by the National Institutes of Health (R01GM113230 to L.P.). We thank Lakshmi Kirkire for keeping this project going after Y.L., Jie Xiang and Patricia Graham for technical advice and suggestions, and Jessica Hernandez for technical assistance.

## Supplementary Information

### Materials and Methods

#### Embryo collection

*O. fasciatus* were reared in 30 x 18 x 20 cm plastic cages. At the beginning of an embryo collection period, a piece of cotton was placed on top of a small section of paper towel and added to each adult *O. fasciatus* cage; the towel barrier ensured that no older embryos already in the cage would be collected. At the end of the collection period, the cotton was removed from the cage and kept at 25°C until embryos reached the appropriate age. Just prior to fixation, the cotton was carefully picked apart, allowing the embryos to fall on to a piece of white paper below. Approximately 100 ul of embryos were then transferred to each microcentrifuge tube.

#### cDNA synthesis and RT-PCR

Embryos were collected every eight hours over a five-day period and maintained at 25°C until reaching the appropriate age. Enough RNA*later* (Invitrogen) was added to cover embryos, and tubes were stored at −20°C until RNA extraction. RNA was extracted using TRIzol (Invitrogen) and the RNeasy Mini Kit (Qiagen). To synthesize cDNA for RT-PCR, RNA from three appropriately aged 0-8 h after egg laying (AEL) collections was pooled for each time point (0-24 h, 24-48 h, 38-72 h, 72-96 h, and 96-120 h) to ensure that a diversity of ages within each time frame was represented. cDNA was prepared using the QuantiTect Reverse Transcription Kit (Qiagen). Primers for *Of-eve* and *Of-E75A* were designed based on previously published sequences (Erezyilmaz et al., 2009; Liu and Kaufman, 2005). Primers for *Of-ftz* and *Of-ftz-f1* were designed based on sequences isolated from cDNA by degenerate PCR, followed by 3’ and 5’ RACE using an RNA ligase-mediated RACE kit (Ambion) or the SMARTer® RACE 5’/3’ Kit (Takara Bio). All other primers were designed based on sequences in the *O. fasciatus* genome. To check for primer specificity, cDNA from the five collections described above was mixed, yielding 0-120 h cDNA. Primers were used to amplify this mixed stage cDNA, and PCR products were sequenced (Genewiz).

#### Embryo fixation, in situ hybridization, and SYTOX staining

Embryos were fixed as described by Liu and Kaufman (2004) with modifications made by Ben-David and Chipman (2010) and by our lab. Briefly, tubes containing ~ 100 ul embryos in ~600 ul water were briefly and gently spun down until submerged. Tubes of submerged embryos were placed in a boiling water bath for 3 min, then chilled on ice for 6 min. Tubes were then laid on their sides and shaken for 20 min at 250 rpm in 1:1 heptane:12% paraformaldehyde. The paraformaldehyde was then removed and replaced with 600 uL methanol. Tubes were then shaken by hand for about 30 s. Heptane and methanol were removed, and embryos were washed three times in methanol. At this point, we sometimes stored the embryos in methanol and completed the remaining steps on the day of use; otherwise, all steps were performed in one day. Embryos were transferred stepwise from methanol to PBST, and eggshells were manually removed. Embryos were washed in 4% paraformaldehyde/PBST for 1.5 h and used immediately or stored in methanol at −20°C.

Antisense RNA probes were synthesized using digoxigenin or fluorescein RNA labeling mix (Roche) and DNA templates generated by PCR amplification of staged cDNA with reverse primers containing the T7 promoter sequence. The T7 RNA transcription reaction was carried out at 37°C for 2h, followed by precipitation in cold ethanol and LiCl. Since our original probes were designed to hybridize to conserved domains, we designed additional probes for *odd, prd, run*, and *slp* that were gene-specific to ensure that our results were not obscured by cross-hybridization with related genes. Gene-specificity was confirmed by conducting a BLAT search of the *O. fasciatus* genome using the probe sequence as the query; if only the sequence of the target gene matched the probe sequence, we considered the probe gene-specific.

In situ hybridization was carried out as previously described either by Ben-David and Chipman (2010) or Liu & Kaufman (2009), with the addition of a 1 h 5% BSA blocking step prior to blocking with 10% sheep serum. Either BCIP/NBT or BCIP/INT was used as the chromogenic substrate for alkaline phosphatase.

Nuclear stains were carried out by rocking fixed embryos in 1:1000 SYTOX Green (Invitrogen) in PBST for 30 min at room temperature. Embryos were then washed three times in PBST.

#### Imaging

Blastoderm-stage embryos were photographed in PBST. In blastoderm-stage embryos, the anterior pole is somewhat tapered; this morphological feature was used to orient embryos anterior-left. Otherwise, germbands were dissected out, transferred to glycerol, and mounted on a slide. Imaging was done with an AxioCam MRc camera (Zeiss) mounted on a SteREO Discovery.V12 dissecting microscope (Zeiss) or Axio Imager.M1 compound microscope with DIC (Zeiss). When necessary, image stitching was performed in Fiji using the pairwise stitching plug-in (Preibisch et al., 2009) or Adobe Photoshop.

#### Isolation of *Oncopeltus* orthologs of *Drosophila* pair-rule genes

To isolate candidate orthologs of *D. melanogaster* pair-rule genes (PRGs), the *O. fasciatus* draft genome was searched using *D. melanogaster* amino acid sequences as queries. All scaffolds that matched with high significance to the query sequence were investigated (e-values usually on the order of 1e-10 or less). Once several gene candidates were identified, multiple sequence alignments, reciprocal BLAST searches, and phylogenetic analyses were conducted to identify the most likely ortholog. Full or partial gene sequences were then isolated by PCR and confirmed by sequencing. In some cases, 3’ RACE was also performed. The gene structures are summarized in Figure S1.

For *odd-skipped (odd)*, three candidate orthologs were found in the genome. Conceptual translation yielded highly conserved zinc finger DNA binding regions for all three which were nearly identical to each other, 94% identical to that of *Tcas-odd*, and 87% identical that of *Dmel-odd* (Fig. S2A). In *Drosophila*, *odd* has two paralogs, *brother of odd with entrails limited* (*bowl*) and *sister of odd and bowl* (*sob*) (Hart et al., 1996). To distinguish between these three closely related genes, BLAST searches were used to gather *odd*, *sob*, and *bowl* sequences from a variety of insect species and outgroups. Deduced amino acid sequence motifs helped to identify the ortholog of *sob* on scaffold 208 and *bowl* on 3993 and 1496; phylogenetic analysis supported these designations (Fig. S2A). The candidate sequence on scaffold 1311 formed a clade with *odd* sequences from the hemipteran *H. halys.* Thus this candidate was designated *Of-odd.* Others have independently assigned the same identities to these sequences (Panfilio et al, 2017). 3’ RACE was performed to isolate the 3’ UTR. A ~900 bp 3’ UTR isolated, only about half of which was found on scaffold 1311; the rest was found on scaffold 7300.

For *odd-paired (opa)*, only two scaffolds matched the query sequence with high significance. The sequence on scaffold 2669 extended approximately halfway through the highly conserved zinc finger domain of *opa* orthologs. The sequence on scaffold 4736 continued this sequence beyond the end of the zinc finger domain. The candidate produced by merging the sequences on these two scaffolds contains a conserved region of zinc fingers that match the *Dm-*Opa zinc fingers with 81% amino acid identity; thus we designated this partial gene sequence *Of-opa* (Fig. S1B).

*paired (prd)* orthologs encode a paired box and homeobox, but lack the octapeptide motif shared by closely related Pax3/7 family members Gooseberry (Gsb) and Gooseberry-neuro (Gsb-n) (Keller et al., 2010). Two scaffolds could be merged to complete a sequence with a *prd* domain, homeodomain, and octapeptide motif NHSIDGILG, which matches the canonical Gsb octapeptide motif (Keller et al., 2010). This candidate from merged scaffolds 2098 and 279 was designated as *Of-gsb*, in agreement with (Ginzburg et al., 2017). Merging two other scaffolds (1714 and 867) containing *prd* candidate sequences yielded a sequence with a *prd* box, homeobox, and lacking an octapeptide motif, designated as *Of-prd*. The *prd* domain encoded by *Of-prd* shares 86% overall amino acid sequence identity with that of *Dm-*Prd, with 93% identity in the homeodomain. After PCR was performed to verify the sequence, two major products were obtained. By sequencing, these were found to be two apparent isoforms of *Of-prd*, one with a 75 bp insertion within the *paired* box. Comparison with other hemipteran *prd* sequences revealed that insertions of variable length at this locus appear to be fairly common. 3’ RACE was performed to extend the known sequence through the 3’ UTR.

*Drosophila* have four Runx family members, all of which share a conserved Runt domain and a C-terminal VWRPY motif, necessary for interactions with the corepressor Groucho (Aronson et al., 1997; Bao and Friedrich, 2008). Searches for *runt (run)* in the *O. fasciatus* genome yielded four candidate sequences containing Runt domains, likely orthologs of *run, lozenge, runxA*, and *runxB*. Two of these genes were found on the same scaffold (scaffold 536), with ~83 kb between them. To determine which of these sequences is the most likely ortholog of *Dmel-run*, phylogenetic analysis using Runt family orthologs from a variety of arthropod species was carried out (Fig. S2D). One candidate sequence (scaffold 536) groups with other *runt* orthologs, contains the Groucho-interacting motif, and was designates *Of-run*. The other sequence on scaffold 536 groups with other *runxA* orthologs; indeed, linkage between *run* and *runxA* is consistent with previous findings (Duncan et al., 2008). The candidate sequence on scaffold 758 was determined to be *lz*, and that on scaffolds 9348 and 2955 is most likely *runxB*.

Similar to *runt*, *hairy (h)* orthologs encode a C-terminal motif (WRPW) which is necessary for interaction with Groucho (Jiménez et al., 1997). Searches in the *O. fasciatus* genome yielded two candidate sequences, both of which contain WRPW motifs and highly conserved Hairy-Orange DNA-binding domains. After phylogenetic analysis with other *h* and related gene *deadpan* orthologs, the candidate sequence from scaffold 1096 was designated as *Of-h*, and the sequence on scaffold 786 was designated *Of-dpn* (Fig. S2E). The hairy domain encoded by *Of-h* is 68% identical to that encoded by *Dmel-h*.

*sloppy paired (slp)* is part of the FoxG subfamily of a fairly large, highly conserved family of *forkhead box* (*Fox*) genes (Grossniklaus et al., 1992; Lee and Frasch, 2004). Searches for *sloppy paired* (*slp)* in the *O. fasciatus* genome yielded eight matches with high significance values (e-value < 1e-20) containing forkhead domains. Phylogenetic analysis was carried out with other *slp* orthologs to determine the most likely *Of-slp* sequence. The candidate sequence from scaffold 497 grouped with other *slp* orthologs and was designated *Of-slp* (Fig. S2F). This sequence matches the partial *slp* sequence found by previously by Liu and Patel (2010) and the deduced amino acid sequence shares 65% and 56% identity in the forkhead domain with *Dmel-*Slp1 and *Dmel-*Slp2.

*Of-ftz-f1, Of-ftz-A*, and *Of-ftz-B* were isolated before the *O. fasciatus* genome sequence was available by degenerate PCR and 3’ and 5’ RACE. For *Of-ftz-f1*, a 2081 bp sequence was isolated, mapping to scaffolds 191, 13634, and 848, and encoding a sequence 564 amino acids long. The DNA binding domain encoded by *Of-ftz-f1* is 97% identical to that encoded by *Dm-ftz-f1*; the ligand binding domains encoded by these two genes are 75 identical.

**Table S1.**
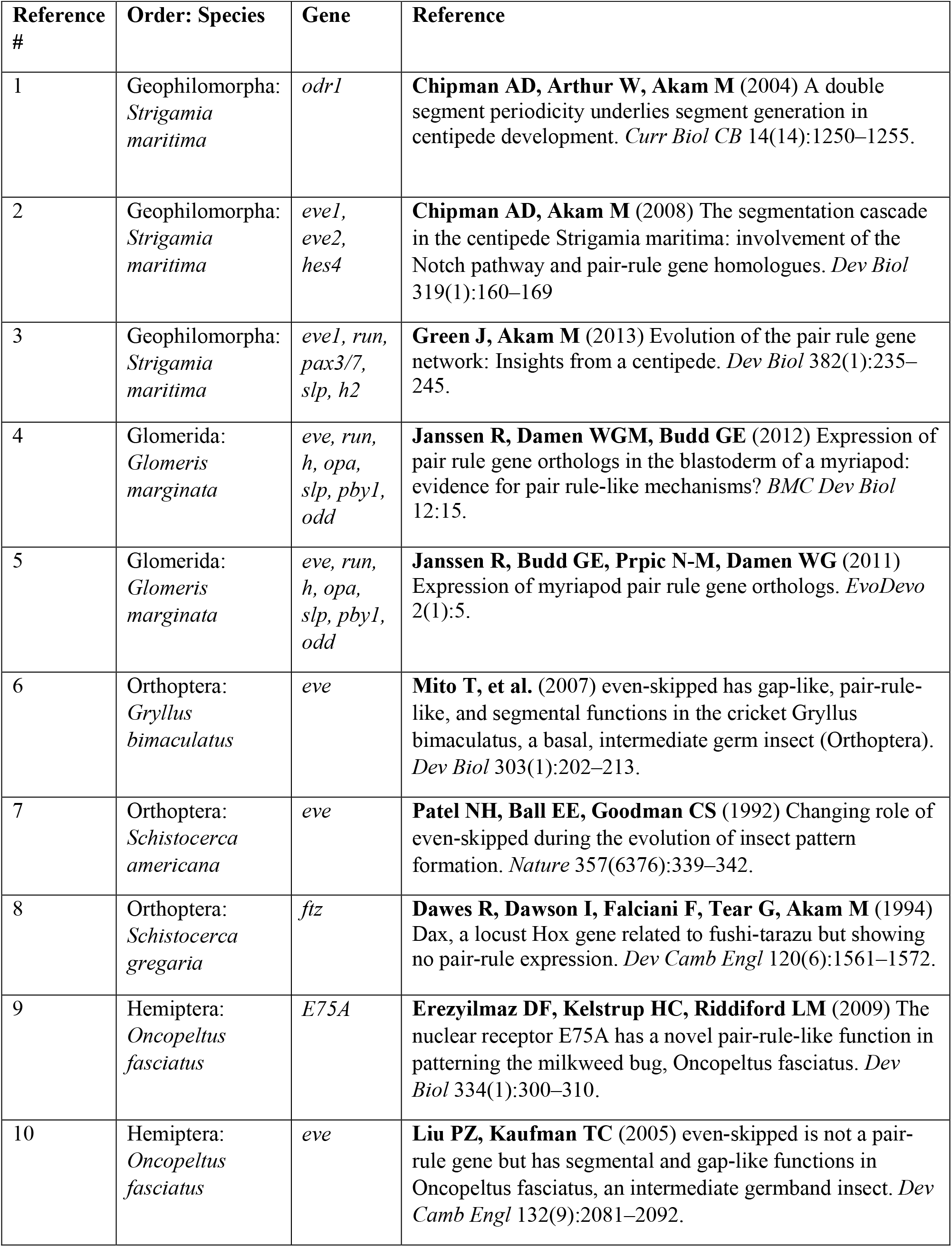
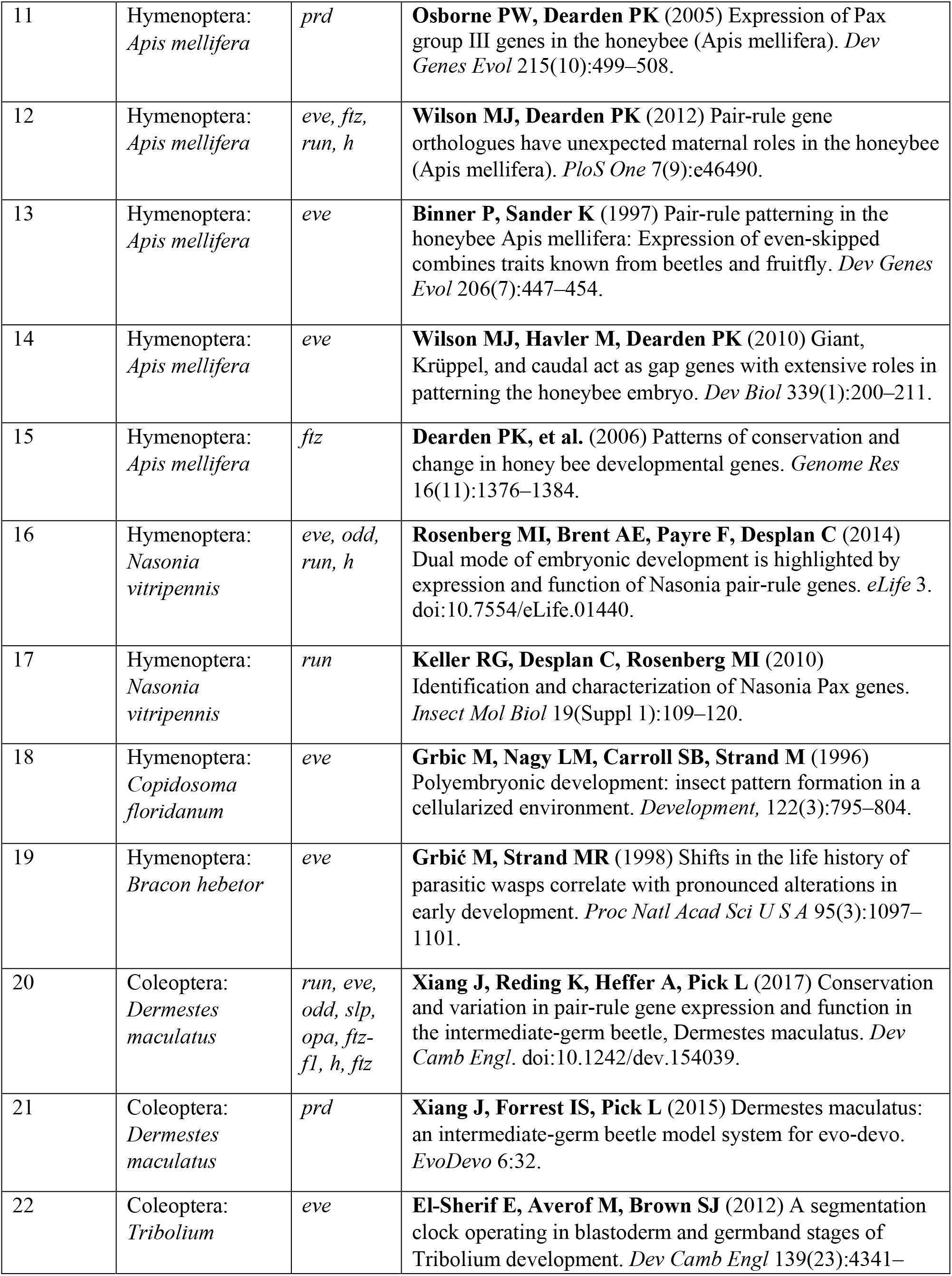
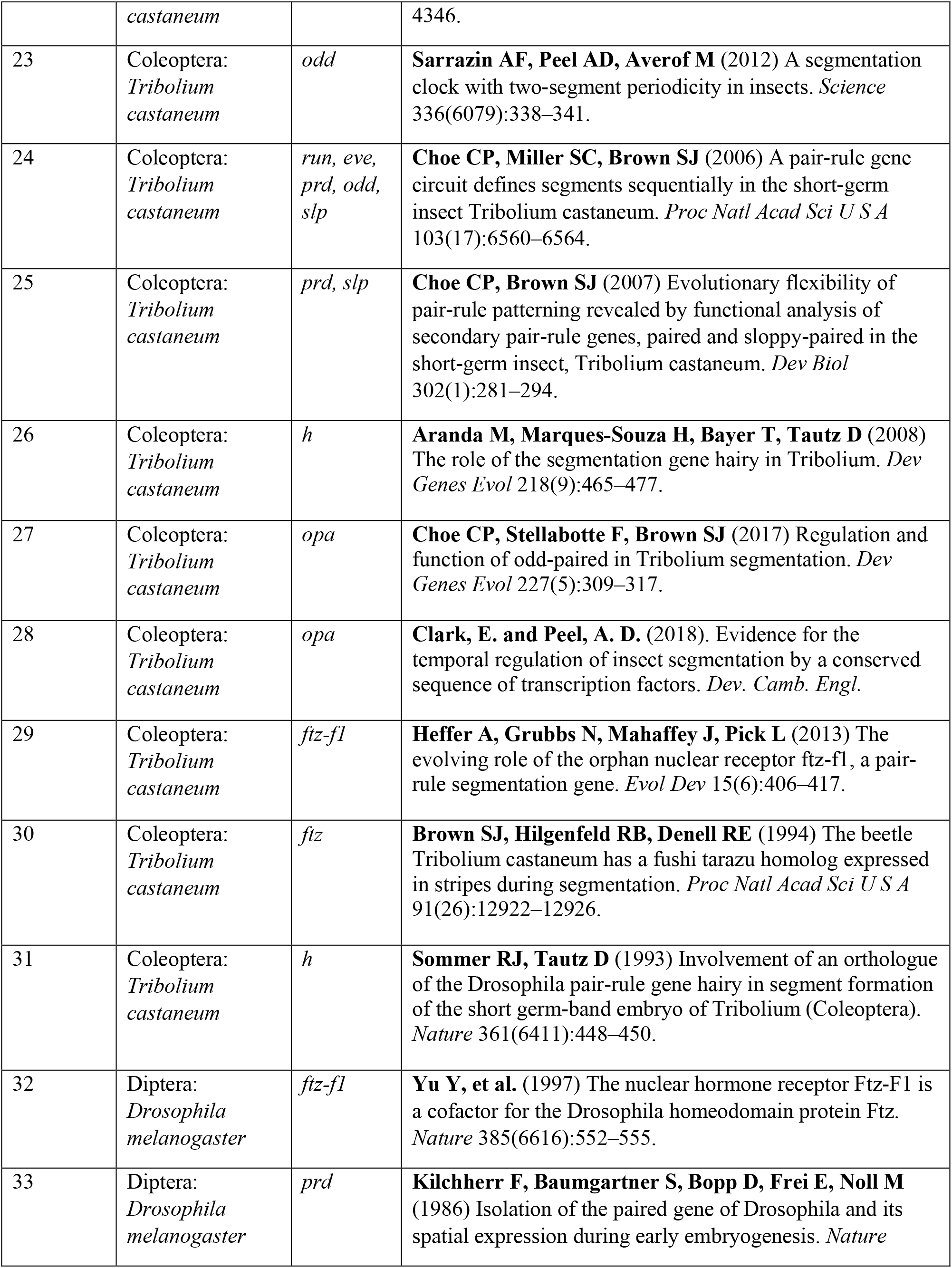
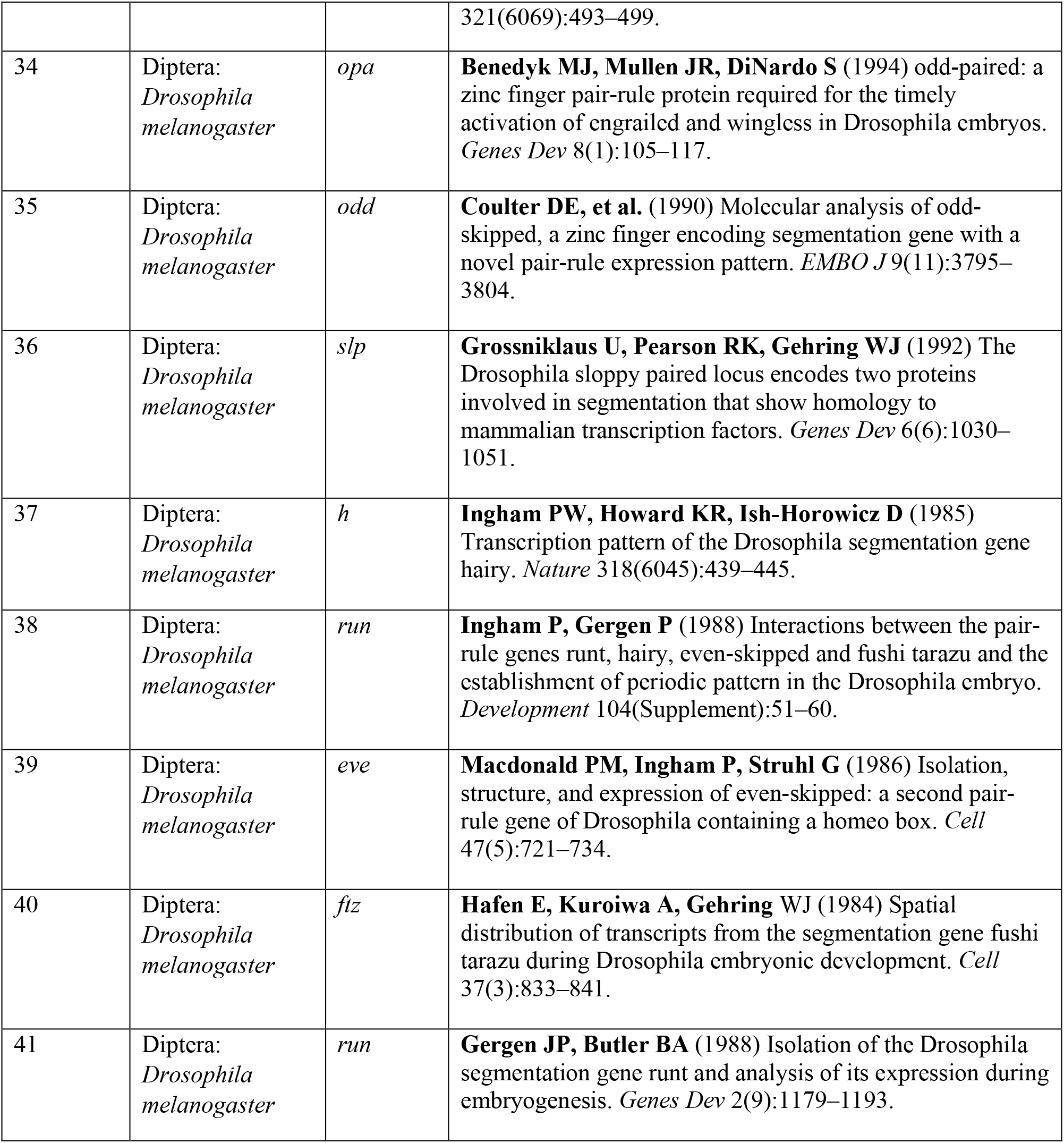
References for PRG literature cited in Fig. 1.

**Table S2.**
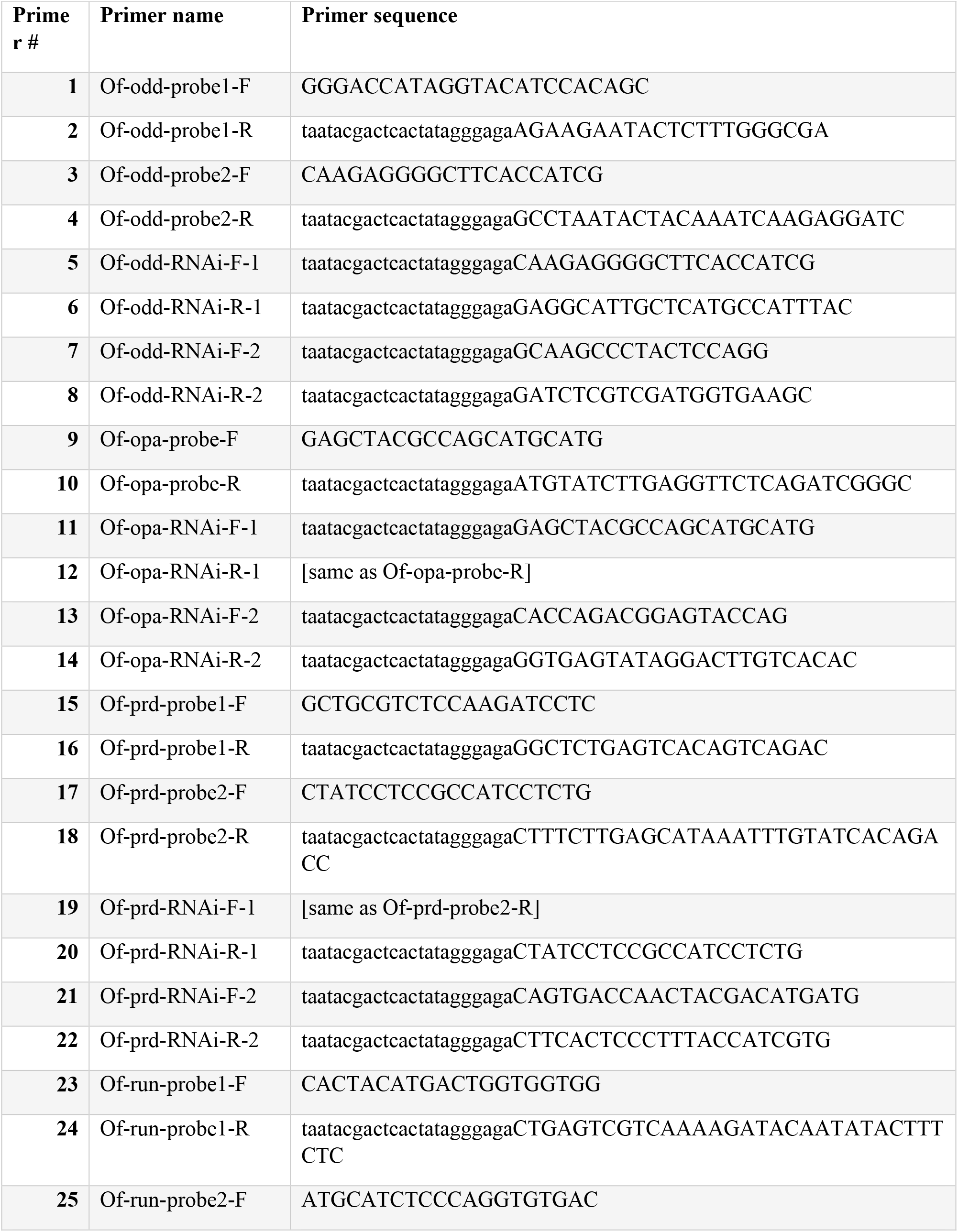
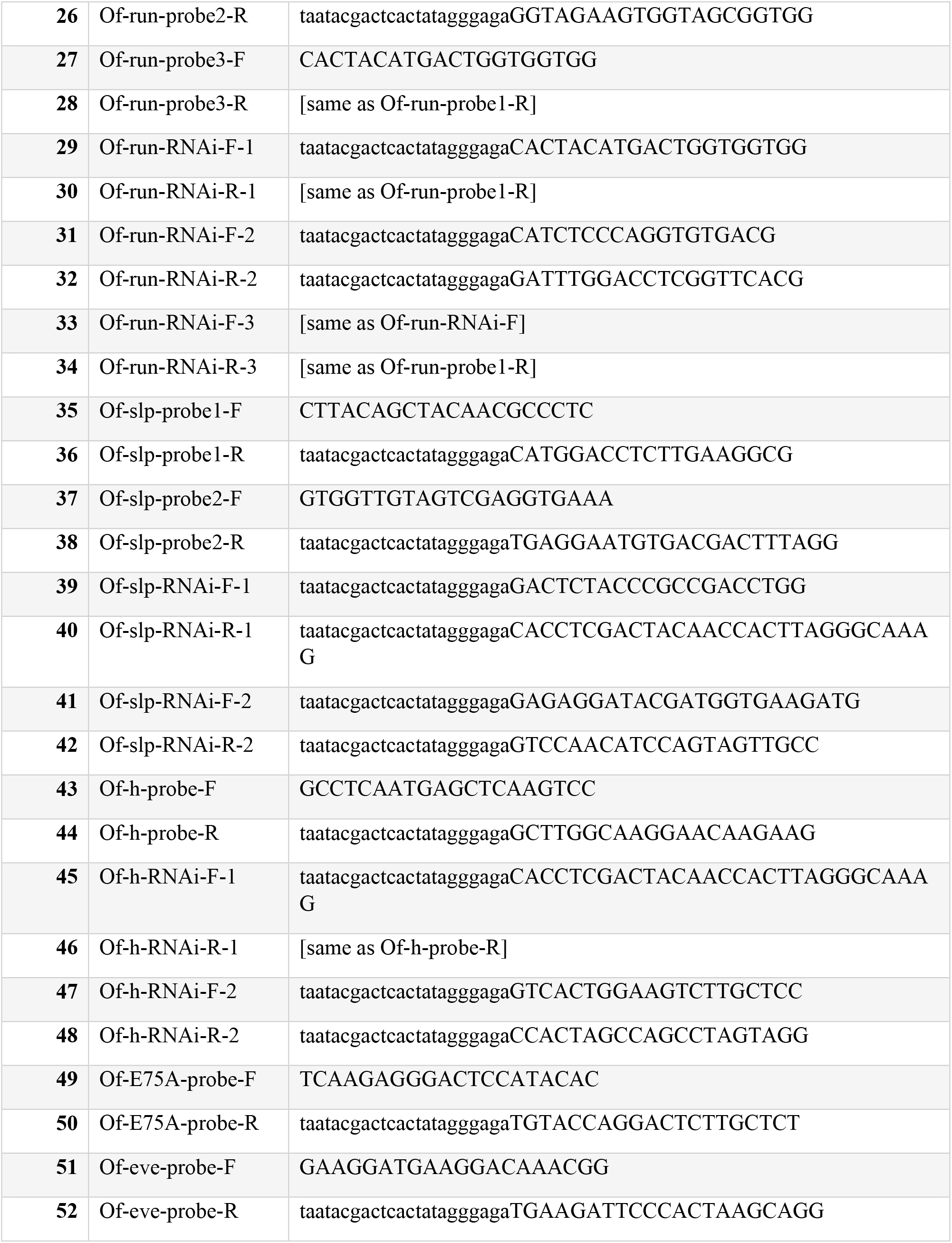
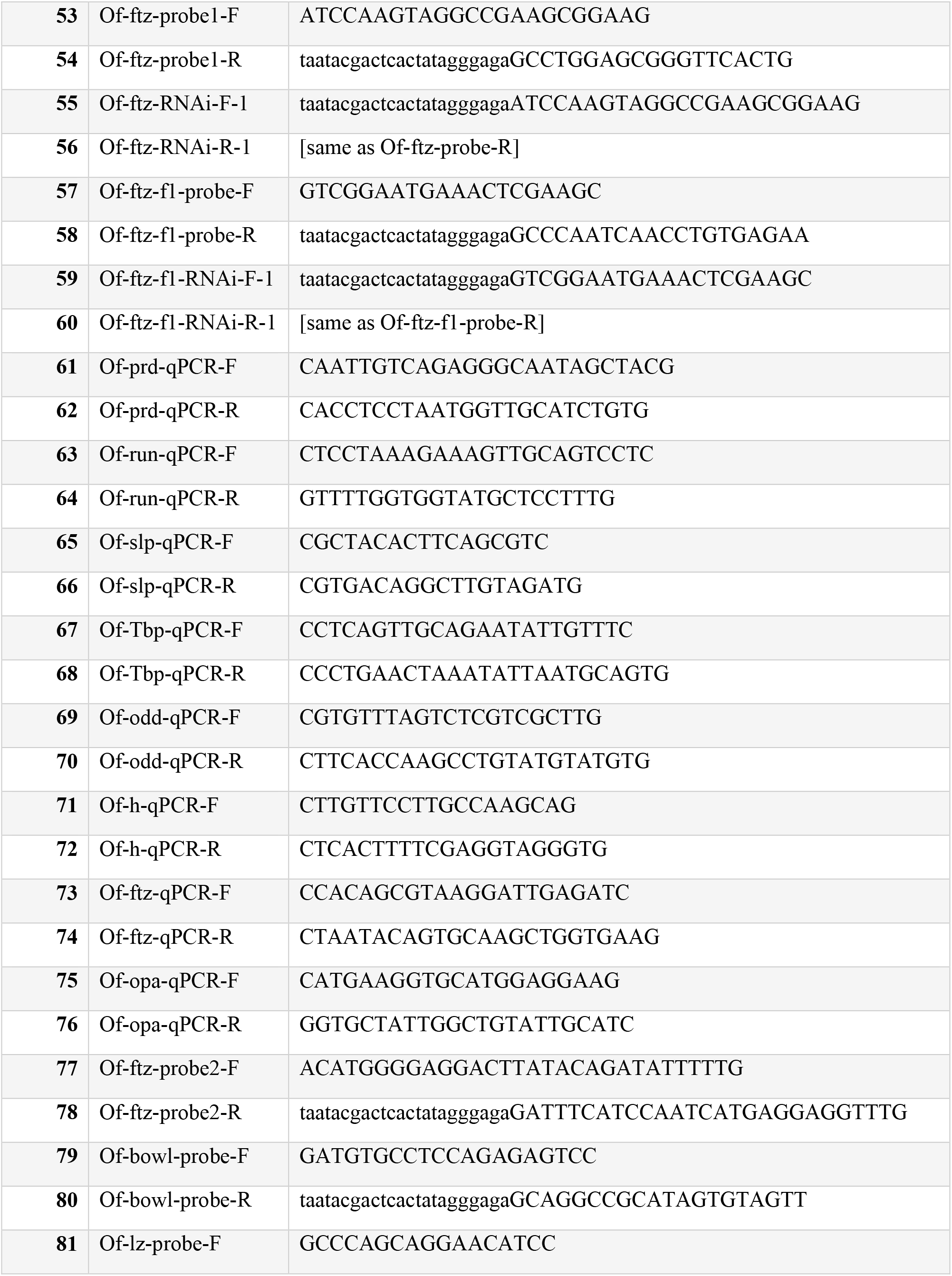
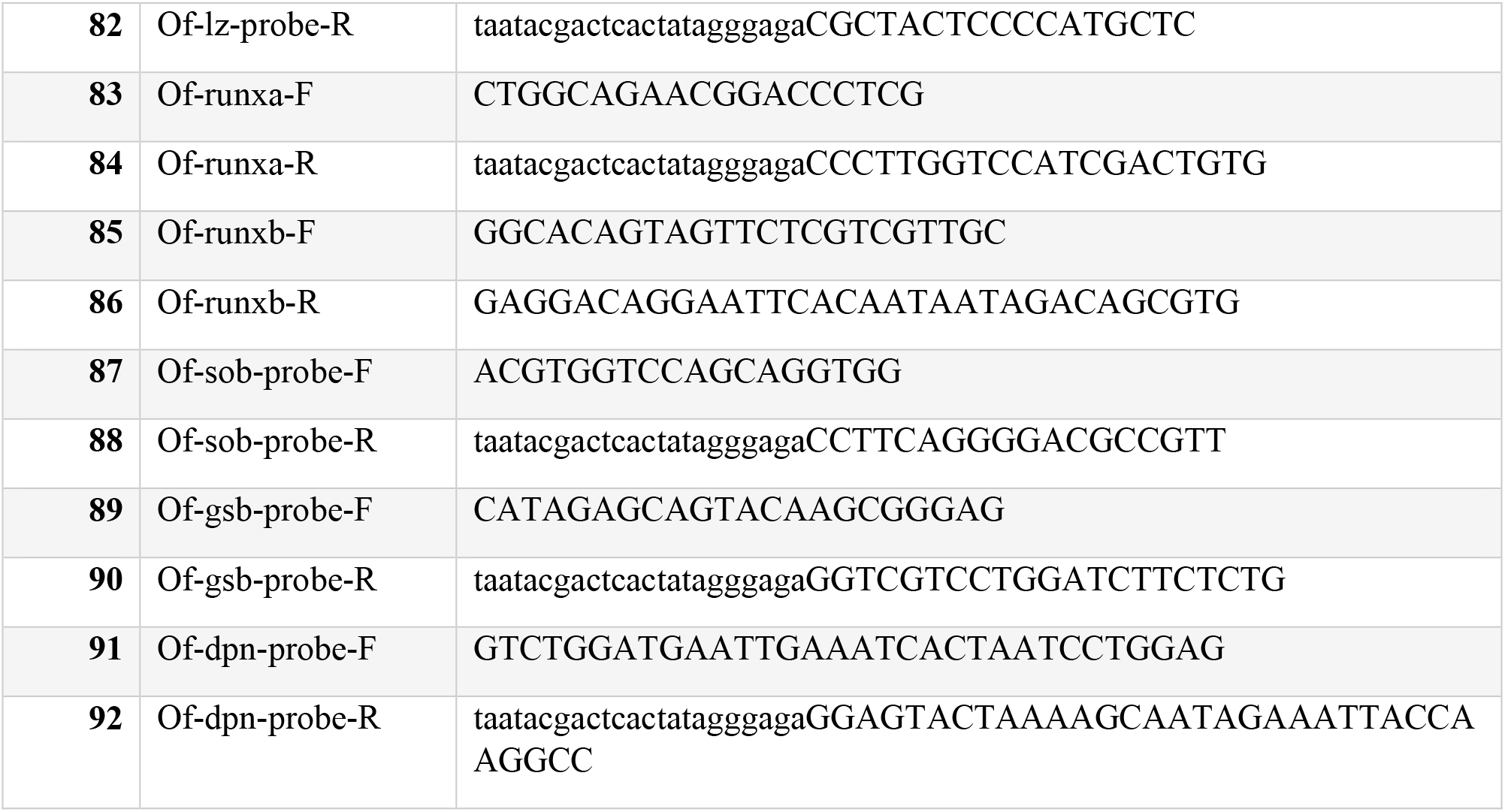
Primer sequences. Lower case letters indicate T7 promoter sequences. The primer numbers correspond to those listed in Fig. S1.

**Fig. S1.**
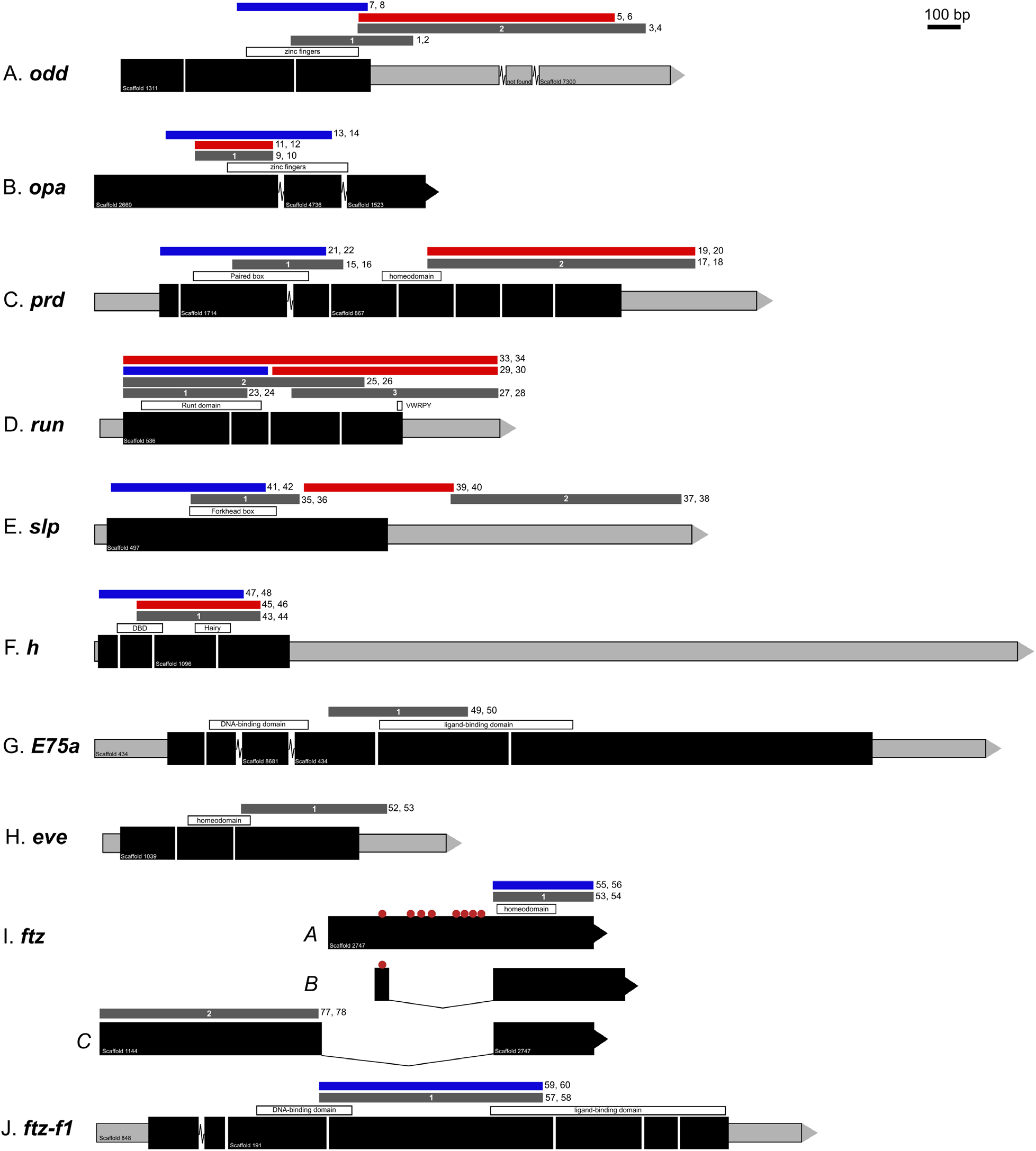
*Of*-PRG-ortholog gene structures. Schematic drawings of the genes analyzed in this study. A) Partial *odd* structure including 3’ UTR; B) partial *opa* structure; C) full *prd* structure; D) full *run* structure; E) full *slp* structure; F) complete *h* structure; G) full *E75A* structure; H) full *eve* structure; I) sequences A, B, and C of *ftz*; J) full *ftz-f1* structure. Black rectangles indicate coding regions. Gray bars indicate UTR. Arrows designate the 3’ end of the transcript. Spaces between rectangles indicate a splice site. Jagged lines between rectangles indicate a scaffold break in the genome. Labeled white bars above structures show the location of signature domains, as indicated. Dark gray bars above structures indicate different probes used for in situ hybridization. Blue bars are regions that were used as dsRNA targets for phenotypic analysis. Red bars indicate other dsRNA targets tested. Red circles in *Of-ftz-A* and *–B* indicate premature stop codons.

**Figure S2.**
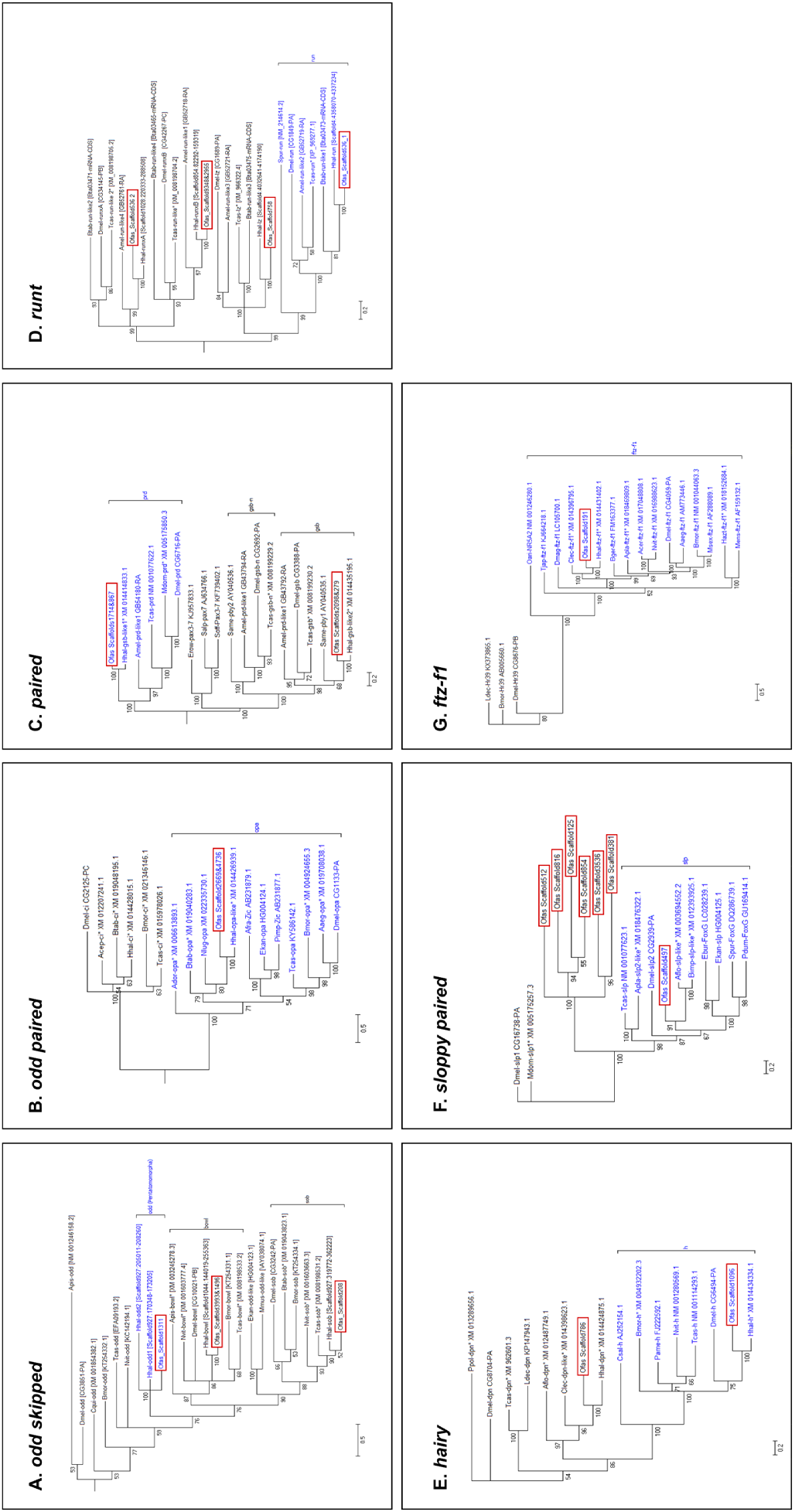
Phylogenetic analysis of gene families was used to designate orthology. Accession numbers are listed next to each gene. A) Three candidate *Of*-*odd* sequence were compared to *odd, sob*, and *bowl* orthologs, B) one candidate *Of*-*opa* sequence was compared to *opa* and *cubitus interruptus (ci)* sequences, C) two candidate *Of*-*prd* sequences were compared to *prd, gsb*, and *gsb-n* orthologs, D) two candidate *h* sequences compared to *h* and *deadpan (dpn)* orthologs, E) seven candidate *Of*-*slp* sequences were compared to several *slp* orthologs, F) one candidate *ftz-f1* was compared to *ftz-f1* orthologs and some *Hr39* sequences. Values at nodes indicate statistical support by posterior probability.

**Figure S3.**
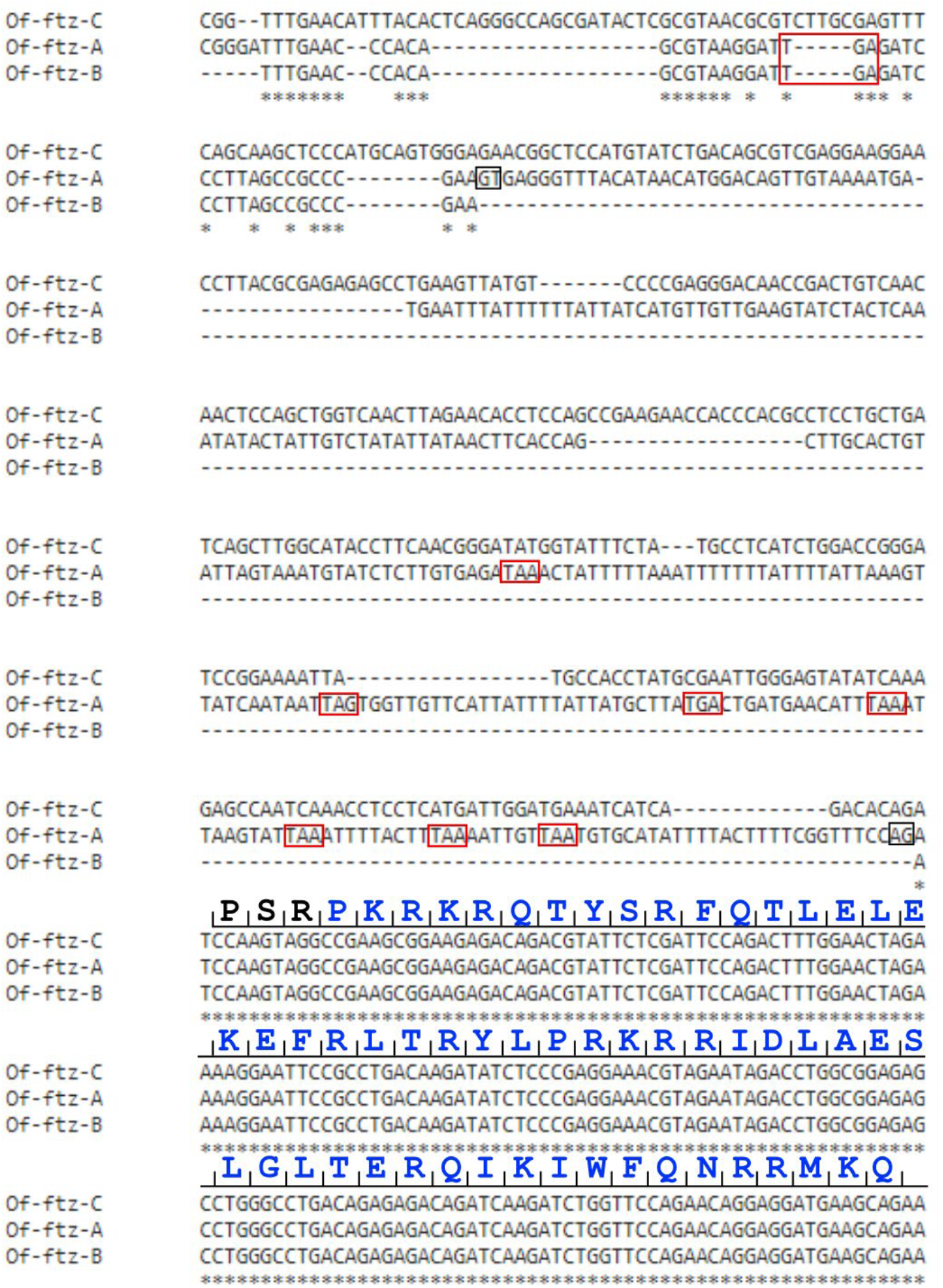
Three *Of-ftz* sequences encode a full-length homeodomain. A partial alignment of *Of-ftz-A*, *-B*, and *-C* nucleotide sequences. Amino acid sequence of the homeodomain is shown in blue. Red boxes highlight stop codons. Black boxes indicate putative donor and acceptor splice sites.

**Figure S4.**
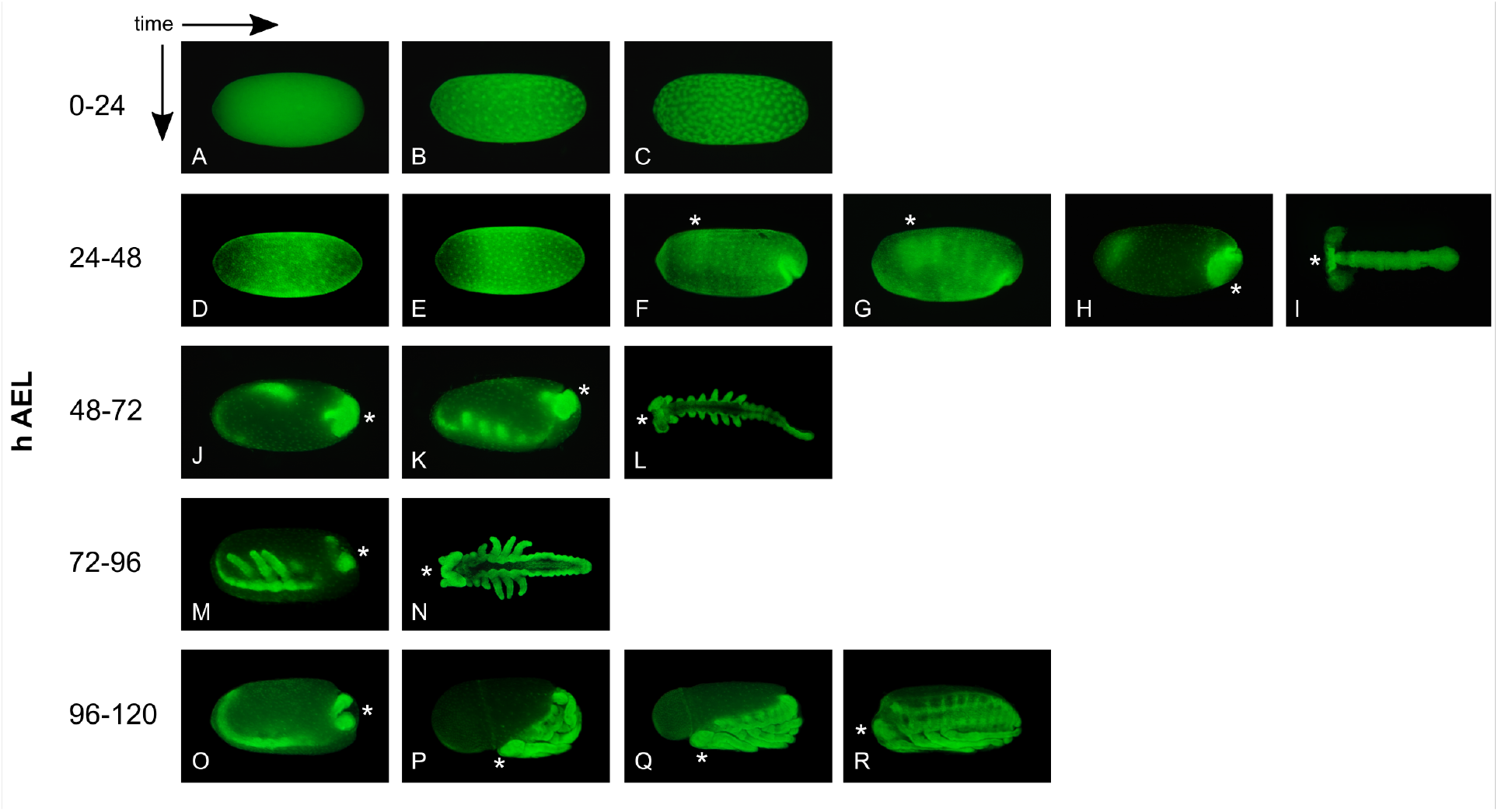
Visualizing *O. fasciatus* development 0-120 h AEL. Embryos were stained with the nuclear stain SYTOX Green; an asterisk shows the location of the head region throughout the embryo’s development; A-C) 0-24 h AEL. Nuclei at the center of *O. fasciatus* embryos multiply and migrate to the surface of the embryo, where they cellularize forming a blastoderm. D-I) 24-48 h AEL, germband in I was dissected from embryo shown in H. The blastoderm cells move toward the posterior pole of the embryo as an invagination pore forms. Anatrepsis continues as the germband grows through the yolk along the ventral side of the embryo from the posterior toward anterior pole. Although the embryo undergoes major movements during its development, the term “anterior” will be defined as the end at which the head is located at the time of hatching. J-L) 48-72 h AEL, germband in L was dissected from embryo shown in K. Appendage primordia (antennae through T3 legs) become apparent and the remaining abdominal segments are added sequentially as the germband curls upward and extends toward the posterior pole, nearly reaching the head lobes. M, N) 72-96 h AEL, germband in N was dissected from embryo shown in M. Germband retraction. O-R) 96-120 h AEL. Katatrepsis occurs followed by dorsal closure.

**Figure S5.**
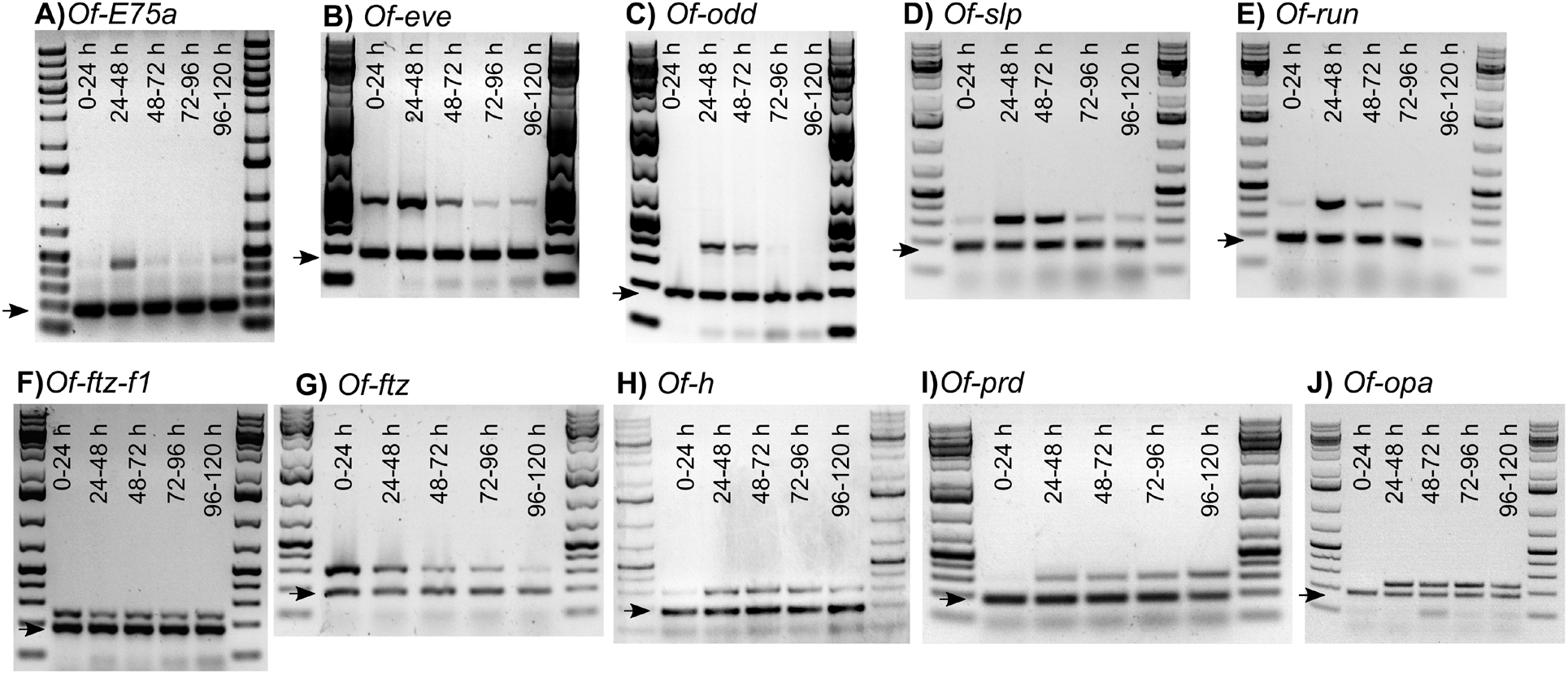
All *Of*-PRG orthologs are expressed during early embryonic development. Time of expression was determined by RT-PCR for A) *Of-E75A*, B) *Of-eve*, C) *Of-odd*, D) *Of-slp*, E) *Of-run*, F) *Of-ftz-f1*, G) *Of-ftz*, H) *Of-h*, I) *Of-prd*, and J) *Of-opa*. RNA was collected from embryos aged to: lane 1) 0-24 h AEL, 2) 24-48 h AEL, 3) 48-72 h AEL, 4) 72-96 h AEL, and 5) 96-120 h AEL. These stages align with those presented in Fig. S4. *actin* primers were included in each reaction, amplifying a ~200 bp product (arrows) as an internal positive control.

**Fig. S6.**
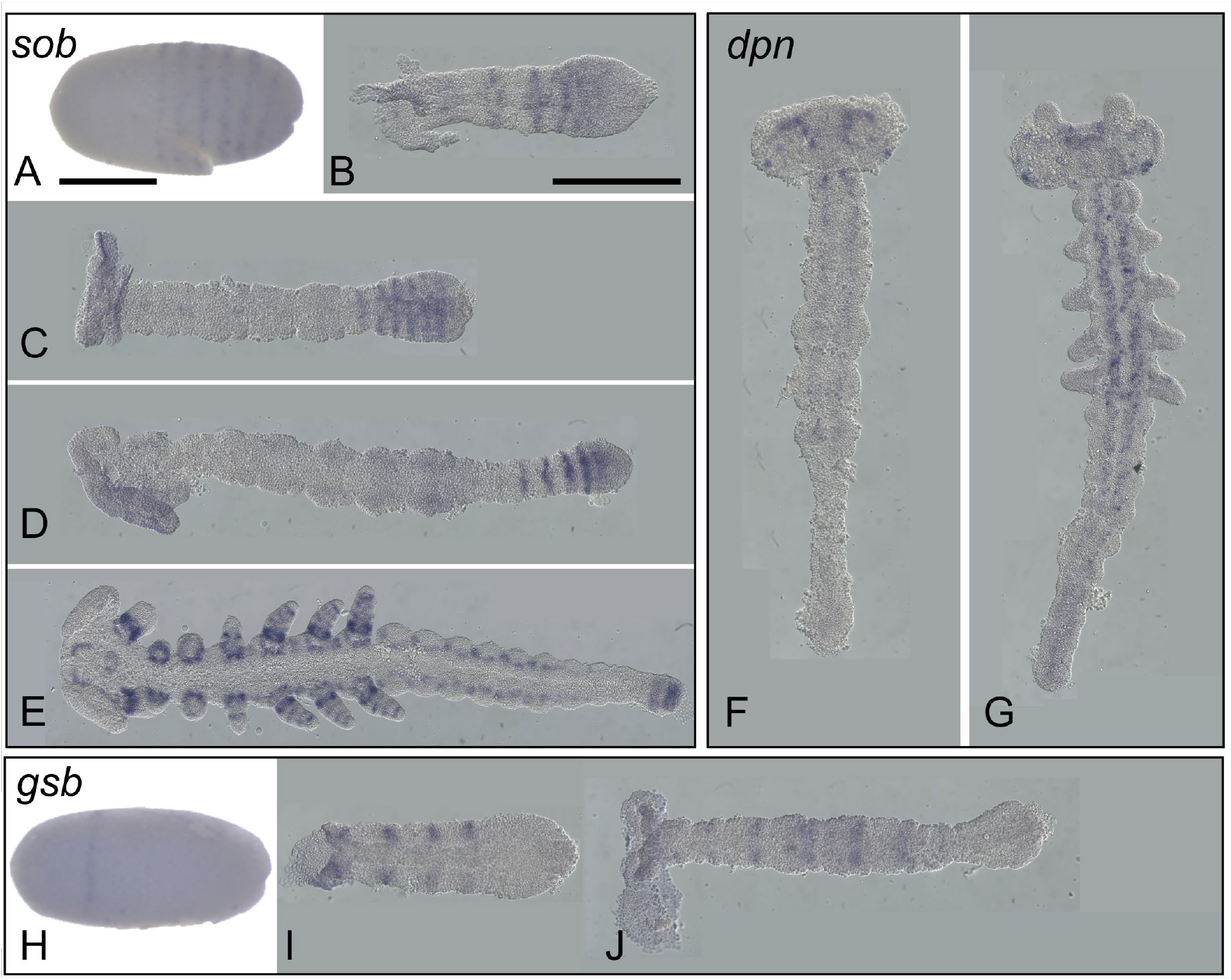
*Of-*PRG paralogs are not expressed in pair-rule-like patterns. A-E) *Of-sob* is expressed in a pattern which appears to be identical to *Of-odd*, in six segmental stripes at germband invagination (A), and in the anterior SAZ throughout germband elongation (B-D). In later-stage germbands, *Of-sob* is expressed in stripes in the appendages, in dots along the lateral sides of the abdomen, and at the posterior terminus (E). F-G) *Of-dpn* is expressed in the head lobes in early germbands (F) and along the midline in later germbands (G). H-J) *Of-gsb* is expressed in one stripe in the anterior of blastoderm-stage embryos at germband invagination (H) and segmentally during germband elongation (I-J). Scale bars correspond to approximately 0.5 mm.

**Figure S7.**
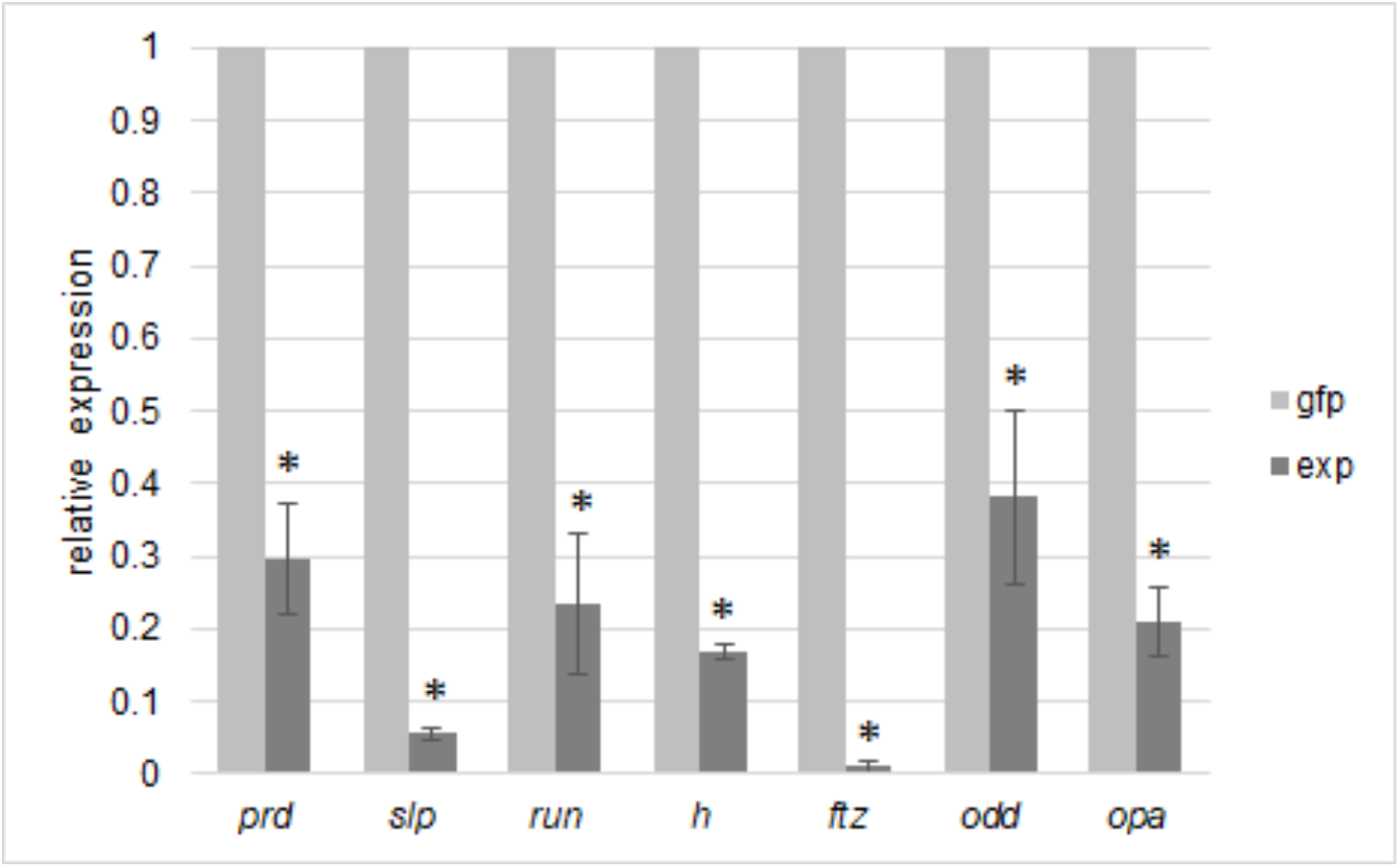
RNAi knockdown is validated by qPCR. Gene expression fold change was calculated relative to a *gfp* control using the 2^−ΔΔCT^ method. *Tbp* was used as the reference gene. Error bars indicate standard error. Statistical significance (α = 0.05) is indicated by an asterisk.

**Fig. S8.**
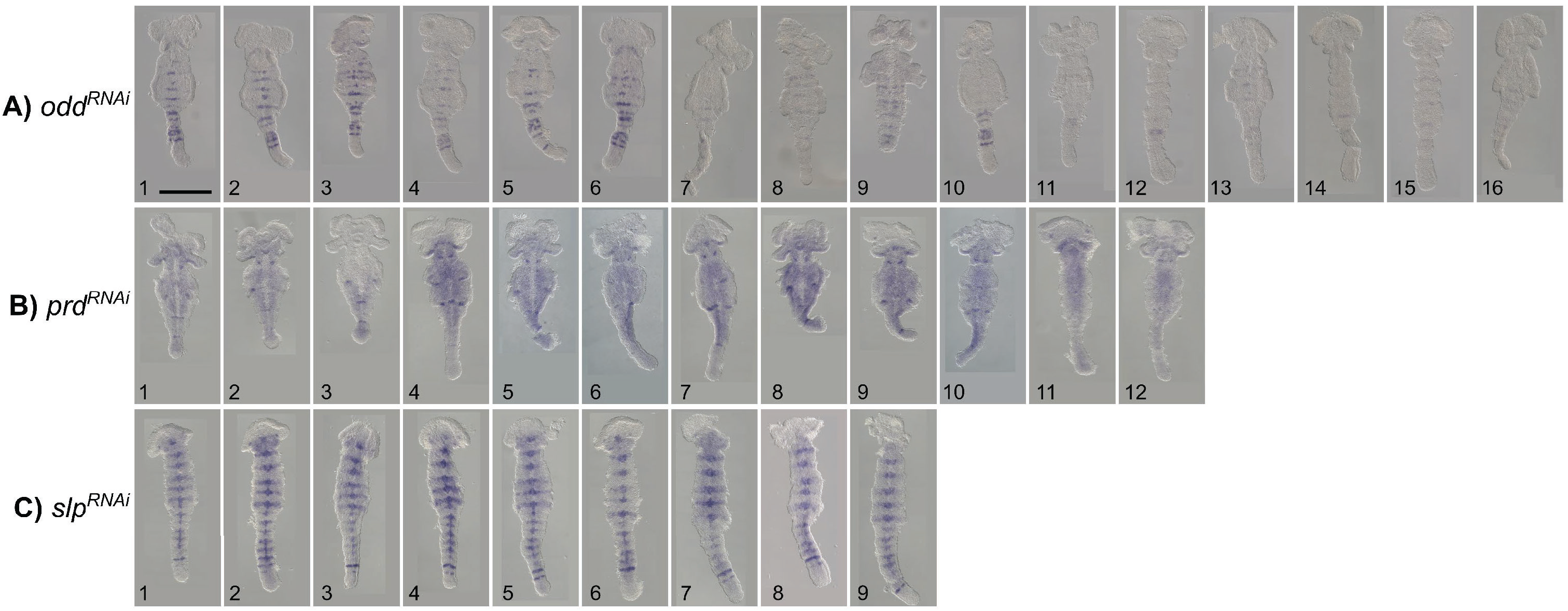
Range of outcomes for PRG ortholog pRNAi. A) *odd* ^pRNAi^ resulted in reduced expression of each *inv* stripe (1-6) or nearly complete loss of each *inv* stripe (7-16) as well as fusion of gnathal and thoracic appendages (9, 16); B) *prd* ^pRNAi^ resulted in nearly complete loss of each *inv* stripe and truncated embryos; C) five *inv* stripes were consistently observed in the gnathal and thoracic regions of *slp* ^pRNAi^ offspring, *inv* expression was expanded especially along the midline of the embryo, and appendages failed to develop. Scale bar corresponds to approximately 0.5 mm.

**Figure S9.**
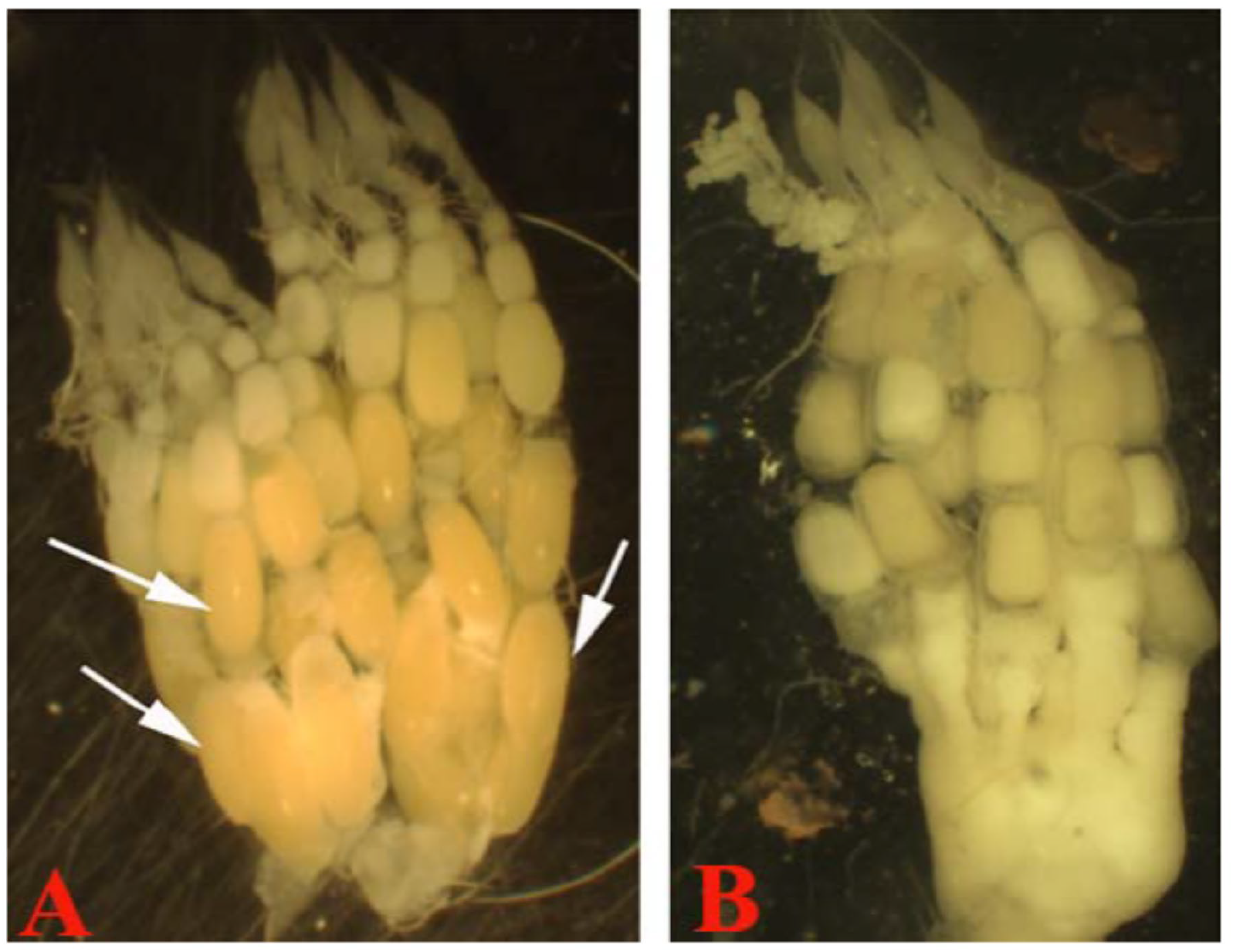
*Of-ftz-f1* ^pRNAi^ disrupts oogenesis. Ovaries from A) a wild-type female, and B) a *ftz-f1* dsRNA-injected female. Wild-type oocytes grow and develop an oval shape and orange color as they mature and move posteriorly. In contrast, the oocytes of *ftz-f1*^RNAi^ ovaries are roughly the same size, with a square-type shape and lighter color, along the length of the ovariole.

**Fig. S10.**
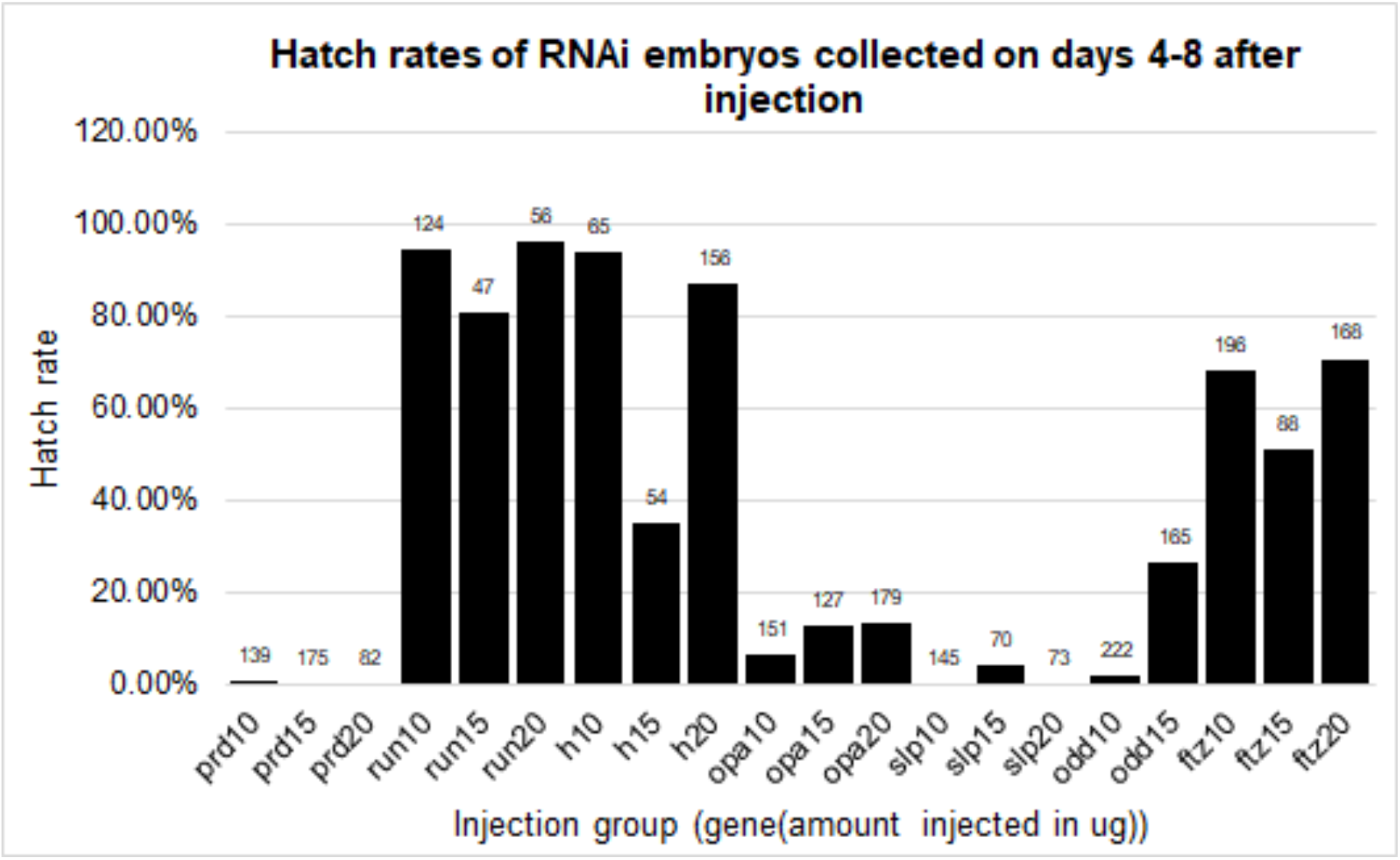
Hatch rates of pRNAi offspring collected 4-8 days after maternal injection. Different amounts of dsRNA, as indicated, were injected into adult females to test the appropriate amount for knockdown of each gene. Hatch rates of embryos collected from each group of females are shown, with the number of embryos analyzed (n) above each bar. Surprisingly little variability in hatch rates was observed for the different amounts tested. Hatch rates were notably suppressed for all amounts of *opa, odd, slp*, and *prd* dsRNA injected.

